# PanTEon: a cross-kingdom framework to guide the design of transposable element classifiers

**DOI:** 10.64898/2026.04.01.715927

**Authors:** Simon Orozco-Arias, Iamil Ferrer-Pomer, Fabiana Rodrigues de Goes, Simon Gaviria-Orrego, Juan Gómiz-Fernández, Jordi Llatser-Torres, Alexandre R. Paschoal, Romain Guyot, Toni Gabaldón

**Author notes:** Corresponding authors: Simon Orozco-Arias; Toni Gabaldón.

## Abstract

Transposable elements (TEs) are major drivers of genome evolution, yet their annotation and classification remain inconsistent and hard to reproduce across species. Fragmented repeats, lineage-specific mutations, and the use of heterogeneous taxonomies across databases and tools make comparative analyses difficult and hamper progress in TE biology. To address this, we developed PanTEon, a cross-kingdom deep learning framework for reproducible TE classification that combines a harmonized database with an open, modular benchmarking platform. The PanTEon Database is an automatically curated, taxonomically broad TE repository spanning animals, plants, and fungi. The PanTEon platform standardizes training, evaluation, and inference across nine machine learning methods, while being extensible to user-defined architectures. We use this framework to benchmark state-of-the-art machine learning-based TE classifiers and find that performance varies markedly across TE superfamilies and major eukaryotic lineages. Ensemble approaches and phylum-specific models improve predictive F1 scores, but cross-species generalization remains challenging. Together, the PanTEon Database and platform provide a reproducible, scalable, and extensible foundation for TE classification, enabling standardized evaluation of future AI methods and supporting community-driven annotation efforts.

## 1. Introduction

Thanks to significant advances in sequencing technologies, it is now easier than ever to generate and analyze complete genomes for a wide range of species. Sequencing companies are continuously increasing the quality and length of sequencing reads while reducing costs (1), thereby improving our capacity to generate more accurate and complete genome assemblies (2). In addition, complementary sequencing and mapping approaches, such as Hi-C, Optical genome mapping (OGM), short-read sequencing, and other orthogonal technologies, are widely employed to improve error correction and scaffolding, further enhancing the accuracy and completeness of genome assemblies (3). Although genome assembly is still a very challenging task, with the right amount of data and mixing different sequencing strategies, it is possible now to generate a high-quality assembly (4, 5). As a result, the so-called “Telomere-to-telomere” (T2T) genome assemblies are now increasingly widespread (6–9), providing unprecedented access to genomic regions that were previously excluded. Similarly, contemporary gene annotation frameworks have achieved high levels of automation and accuracy (10–14). These platforms enable the systematic annotation of genes across thousands of genomes, often with minimal manual intervention, drastically reducing the time and labor previously required, while maintaining high-quality annotation output (15). In contrast, the annotation of other genomic elements remains problematic, particularly those involving highly repetitive sequences like transposable elements(16–18).

Genome sequences are not static, as they entail components that can ’move’ from one location to another. These components, called transposable elements (TEs), can be activated by different external or internal factors, such as radiation, pollution, viral infections, or environmental changes (19, 20), and are closely related to processes of adaptation and the evolution of species (21, 22). Although historically understudied due to both conceptual and technical limitations (23, 24), it is now clear that TEs are crucial genomic elements that play an important role by dynamically shaping genome structure and function (Iwasaki et al., 2025; Prokopov et al., 2025), and acting as sources of cis-regulatory elements. Despite the current widespread awareness of their importance, the study of TEs in most organisms remains limited due to important methodological challenges (27).

TEs possess several characteristics that complicate their study. They are often species-specific; consequently, TEs from different genomes, even from closely related species, can exhibit substantial sequence divergence (20). Furthermore, TEs evolve faster than other genomic elements, resulting in substantial sequence divergence among copies belonging to the same family, ranging from nearly identical to highly degraded elements (28–30). Finally, the generally large number of copies (up to several thousand like Alu or L1 in mammals) and the fact TEs can be inserted within one another (nested insertions), make the accurate detection and annotation of these sequences very complex (31).

The standard workflow for studying TEs consists of four main steps: detection, classification, curation, and annotation (31, 32). Nowadays, a plethora of tools are available that cover all or some of these steps (for an exhaustive list, see TEhub.org (33)). Most existing tools can be classified into the following groups based on their TE prediction strategy: *de novo*, structure-based, homology-based, Artificial Intelligence (AI) based, or a combination of these (34, 35).

Driven by recent advances in genomic sequencing, assembly methods and TE biology, several databases have been constructed to facilitate access to TE consensus sequences detected across thousands of genomes. InpactorDB (36), Repbase (39), Dfam (38), and RepetDB (37) are among the most popular databases. However, the number of curated TEs publicly available in these databases is still very limited. Repbase v.29.08 contained 114,924 TE consensus sequences in 2025 according to a report (68), but payment was required to download the data until recently (June 25^th^, 2026; (69)). Dfam v.4.0 is free but it contains only 30,621 curated TE sequences (according to the Dfam web site accessed on 06/19/2026). Finally, RepetDBv2 v.2.1.3 is composed of 130,033 TE sequences (according to the RepetDB web site accessed on 06/19/2026), which are freely available to download, although only 9,584 have been manually curated (classification and/or structure validated), thereby necessitating several pre-processing steps before use to ensure high quality. The hundreds of thousands of TEs from different species contained in these databases have created a unique opportunity to train AI models for the detection and annotation of these elements. However, this requires facing several challenges associated with this type of data, which is categorical in nature, poorly structured, and extremely large (with thousands of characters per element).

So far, classification has been the task that has benefited most from applications of AI, particularly deep learning (DL). Neural networks such as TERL (40), DeepTE (41), TEClass2 (42), NeuralTE (43), CREATE (34), and BERTE (44), among others, have shown improved classification performance as compared to conventional algorithms. SENMAP (45) is so far the only neural network specifically designed to curate TEs, but it is restricted to a single order (LTR retrotransposons) and for plants. Furthermore, no study has used AI to address the tasks of detection and annotation of TEs. The only implementation of genomic object detection (applicable to the annotation task) is that of YORO (46), which produced promising results for the annotation of internal regions of LTR retrotransposons. Although TEs are attracting increasing attention from algorithm developers, particularly AI experts, the lack of accessible, high-quality and taxonomically broad TE databases, standardized benchmarking methods, performance metrics and a unified classification nomenclature constitute a barrier for the development and use of AI-based algorithms for the prediction and classification of TEs. To address these important gaps, we developed PanTEon, a framework combining two complementary components: i) an open database (PanTEon Database) comprising almost 240,000 automatically curated TEs, across 2,790 species of animals, plants and fungi and from all TE orders, and ii) The PanTEon Platform, a computational framework that enables the simultaneous training and evaluation/benchmarking of multiple AI architectures using a unified dataset, generating a consensus output that facilitates the identification of the most reliable TE classifications across models. In addition, the platform supports TE classification inference using individual or combined models and enables the generation of TE libraries for any taxonomic or TE classification level. We showcase the use of PanTEon Database by benchmarking seven state-of-the-art AI-based classification tools, assessing their taxonomic biases. Finally, we utilized the PanTEon Training Module to re-train and compare the performance of nine machine learning (ML)/DL architectures (the seven used before plus two extra) to predict 30 superfamilies, to generate kingdom/phylum-specific models, and to detect false positives in TE libraries.

## 2. Methodology

### 2.1 Creating the PanTEon Database

The PanTEon Database was designed as a comprehensive and automatically curated collection of TE sequences. The database included 371 fully curated entries from Dfam (38) version 3.9, comprising 351 species and 35 other entries assigned to higher taxonomic levels. We then complemented the dataset with 165 species from Dfam (uncurated sequences), as well as libraries from 12 additional species obtained from APTEdb version 1.0 (RepeatModeler2 outputs only; Pedro et al., 2021). These species were selected because they were not represented among the curated Dfam entries. The chosen libraries were composed of uncurated TE sequences identified by automatic tools, such as RepeatModeler2 (RM2) (Flynn et al., 2020). Because the raw outputs of these tools may contain redundant sequences, false positives, and incomplete or truncated consensus sequences, we processed the libraries with MCHelper (49) to reduce redundancy, remove potential false positives, and extend incomplete TE consensuses. Finally, we only kept those TE sequences that contained all expected features corresponding to their classification. For example, LTR retrotransposons were required to have the two LTRs (at the beginning and at the end), and a minimum of four domains (see table S1 for all requirements).

To further complement the database and increase taxonomic representation, we selected 748 species from the Ensembl 2025 update (50) that were not represented in our database. We downloaded, curated, and filtered the raw RM2 results for these genomes, as described above. Because fungal taxa remain underrepresented in most publicly available TE resources, we further increased representation by incorporating decontaminated genomes from the B-GUT database (51). For these genomes, RM2 was executed using the - LTRStruct parameter to enhance the detection of structurally supported LTR retrotransposons. The resulting raw TE models were subsequently curated following the same standardized pipeline applied to all other species. A complete list of incorporated species, together with their taxonomic classification and TE counts, is provided in Supplementary File 1.

Each species library was processed individually through a standardized workflow that included renaming duplicate identifiers and standardizing classification labels into a consistent hierarchical format (Class / Order / Superfamily; (52)). Superfamily-specific length thresholds were applied as a quality-control procedure to remove sequences likely corresponding to fragmented, chimeric, or erroneously reconstructed TE consensus sequences. Threshold values were determined from a combination of published descriptions of TE structural organization (20, 53–58), and expert-curated TE protocols (35, 59–61). Because TE lengths can vary substantially within some superfamilies, particularly LTR retrotransposons and MITEs, these thresholds were not intended to represent strict biological boundaries. Instead, they were used to exclude extreme outliers that are more likely to arise from annotation artefacts or consensus reconstruction errors. The complete set of thresholds is provided in Table S2. Then, TE sequences (except those originating from curated Dfam) were systematically renamed to guarantee their unique identification within the database, following the format: PDBnumericalID#Classification followed by an empty space and “@Species”. The PanTEon Database and its associated metadata, including source database and original identifiers of each sequence, are available at https://zenodo.org/records/18039746.

### 2.2 Benchmark model selection and installation

Several state-of-the-art methods to classify TEs were chosen based on three criteria: (i) strong empirical validation reported in the literature, (ii) tools based on an ML or DL model, and (iii) an available pre-trained model and software enabling the inference of TE superfamilies. The selected methods were ClassifyTE (62), CREATE (34), DeepTE (41), NeuralTE (43), TEclass2 (42), TERL (40) and Terrier (63).

We installed these methods following instructions provided in their respective GitHub repositories (Table S3), with the only exception that all environments were created and managed using Conda. For tools with multiple pre-trained models, the following were selected: ClassifyTE (combined, trained on Repbase and PGSB (64)), DeepTE (kingdom-specific models for animals, plants, and fungi), NeuralTE (non-TSD model for comparability, as the TSD version requires a genome assembly), TEclass2 (clust_cats_16, default in config.yml), and TERL (DS3, trained on seven databases and classifying 11 superfamilies). The remaining tools offered a single model.

### 2.3 Subset creation and tool execution

Because each classifier was originally developed and trained to predict a distinct set and number of TE superfamilies, direct comparison across methods required harmonization of their classification outputs. To construct the benchmarking datasets, we first identified the TE superfamilies supported by each classifier and determined their intersection across methods. Superfamilies recognized by the largest number of tools were selected for inclusion in the benchmarking analysis (Table S4). Only TE sequences assigned to these selected superfamilies were retained for downstream evaluation. Then, a subsampling was applied to have a balanced dataset across Animalia, Plantae and fungi. Because fungi comprised the smallest dataset (12,914 sequences belonging to benchmarking superfamilies), we randomly selected an equal number of sequences from Animalia and Plantae datasets, ensuring that selected sequences were not Dfam curated sequences (to avoid curation and training bias). Using these TEs, we generated one file per kingdom and one combined file. The classifications provided in these four files were used as the reference labels and the corresponding sequences served as input for each evaluated tool. The primary aim of this benchmark was to assess agreement between the classifications in the PanTEon Database and the predictions generated by the tools, rather than to establish absolute biological correctness. Four tests were executed for all methods, one using the combined input set (TEs from all the kingdoms) and one for each kingdom (Animalia,

Plantae, fungi). DeepTE was the only exception since the tool allows the user to select the kingdom-specific pretrained model. In that specific case, we ran the tool separately on each kingdom-specific input set, and then we merged all the results into one file. All executions were performed with default parameters on an HPC system provided by the Institut Français de Bioinformatique (IFB).

### 2.4 Output standardization and ordering

The output of each tool was converted into a tabular file (.csv) and ordered according to the sequence identifiers in the input sets. In the case of DeepTE, its three separate outputs were first merged into a single file before applying the same procedure.

Superfamily names were then standardized using a synonym dictionary, and the results were consolidated into a single table file containing, for each sequence, the original classification and the prediction of each model at the Class, Order and Superfamily levels (using an in-house script called “metrics_intagrated.py”). This standardized table served as input for the subsequent performance evaluation of individual classifiers and ensemble methods.

### 2.5 Performance evaluation and ensemble classification

For the combined and each kingdom-specific datasets, the performance of each tool was evaluated at the Class, Order and Superfamily levels using the following metrics: accuracy, precision, recall and F1-score (implemented in “metrics_intagrated.py”).

To evaluate the potential enhancement of predictive performance through classifier integration, three ensemble strategies were implemented (through an in-house script called “ensamble_classification.py”): (i) simple majority voting, (ii) weighted majority voting with predefined tool-specific weights, and (iii) a supervised stacking model based on gradient-boosted decision trees (XGBoost). For the weighted voting approach, we defined the following weights favoring the tools with better performance based on the benchmarking results: NeuralTE: 1.5, Terrier: 1.5, CREATE: 1.3, TEclass2: 1.2, DeepTE: 1.1, ClassifyTE: 1.0, and TERL: 1.0. For the stacking approach, the dataset was partitioned into training (60%) and test (40%) sets to train and evaluate the meta-classifier. Each ensemble approach generated a consensus classification for every sequence, and their performance was quantified using the same metrics as before.

To measure tool runtime, we executed each tool 10 times using the subsampled dataset (defined in section 2.3), on the same compute node with identical allocated resources (20 cores and 16 GB RAM). Then, we calculated the average and the standard deviation of the computing time to guarantee a fair comparison.

### 2.6 Benchmark comparison using the Friedman test and Critical Difference diagrams

To compare the performance of multiple TE classification tools across the PanTEon subsampled dataset, we applied the Friedman test followed by the Nemenyi post-hoc procedure, which is a standard approach for evaluating significant differences among classifiers under repeated, matched experimental conditions. For each tool, we computed its average rank across all superfamilies, treating lower ranks as better performance.

These ranks were then used to construct a Critical Difference (CD) diagram, where tools connected by a horizontal bar fall within the CD threshold and therefore do not differ significantly at α = 0.05. The diagram provides an interpretable visual summary of pairwise statistical significance across all methods, highlighting groups of tools with comparable performance and identifying those that are consistently superior or inferior across the benchmark. All analyses were performed using the *SciPy* package and custom visualization scripts implemented in Python (in the in-house script called “compute_f1_retraining.py”).

### 2.7 PanTEon Platform development

Existing tools differ substantially in their approaches to classify TEs, including the use of distinct evaluation strategies (e.g., multiclass (MC) versus hierarchical classification (HC) metrics), training datasets (most commonly derived from Dfam and Repbase), and ML/DL architectures (such as ensemble-based methods, convolutional neural networks, and Transformer models). In addition, these tools adopt heterogeneous nomenclatures and support different numbers of TE superfamilies. Hence, we uniformized all steps while leaving the feature extraction and the ML/DL architecture as reported by each tool. In that way, we could check whether a specific method performs better than the others.

Therefore, we defined the classification problem as a multiclass classification task in which a given AI model receives a DNA sequence and must assign the sequence to one of 30 classes. We defined those 30 superfamilies as the most common superfamilies with at least 100 sequences in the whole dataset. Also, we grouped some superfamilies together following the Dfam classification taxonomy (accessible at https://dfam.org/classification/tree), due to some of the superfamilies having very few samples (Table S5).

We used the same seven tools utilized in the benchmarking section (section 2.2) plus two additional neural networks, Inpactor2_Class (65) and BERTE (44). Then, we developed a Python script based on the ML or DL approach used in each tool, but adapting those that used HC to use MC, to predict 30 labels, and we then integrated it into the PanTEon Platform. The PanTEon Platform was developed in Python version 3.10, using TensorFlow, Keras, and PyTorch as DL frameworks. All the packages were installed using Miniforge and all the “experiments” were executed in the Marenostrum 5 HPC supercomputer at the Barcelona Supercomputing Center. The plant data coming from APTEdb were processed using the Bioinformatics and Pattern Recognition group server (BIOINFO-CP) and the multi-user cluster, both from UTFPR Brazil. PanTEon and all the in-house scripts used in this study are available at GitHub: https://github.com/simonorozcoarias/PanTEon

### 2.8 Re-training and ML/DL architecture benchmarking

We used the PanTEon Training Module for training the nine ML/DL architectures available using the entire PanTEon Database. We split the dataset into training (80%), validation (10%) and test (10%) sets (except for TEclass2, which the split was75% training, 15% validation and 10% testing, following the recommendation provided by the developers), and calculating performance metrics using the test dataset.

We measured runtimes for each architecture in two main processes: feature extraction and model training. For doing that, we trained each ML/DL architecture 10 times using the entire training dataset, in the same compute node with 20 cores and one NVIDIA H100 GPU with 64 GB VRAM. Then, we calculated and reported the average and the standard deviation.

We next evaluated the generalization capability of the models using a curated set of TE sequences from species not represented in the PanTEon database. To this end, we downloaded the complete curated dataset from Dfam v4.0 and retained only sequences from species absent from the PanTEon dataset, resulting in 6,218 sequences. We then removed all non-TE sequences (e.g., satellites and simple repeats). For LTR retrotransposons, the LTR and internal regions were merged based on their sequence identifiers to reconstruct complete elements whenever possible, yielding a final dataset of 2,309 TE sequences. Finally, we generated two evaluation datasets using an in-house script, TE_complete_classifier.py: one containing only complete TEs with the expected structural features (867 sequences; see Table S1 for the complete set of requirements) and another consisting of incomplete or degraded elements (1,442 sequences). The selected sequences encompass seven phyla and 48 species and seven TE orders and 26 superfamilies. These datasets were analyzed using the PanTEon inference module to predict TE classifications, and the PanTEon evaluation module was subsequently used to calculate performance metrics by comparing the predictions against the Dfam-derived reference annotations.

### 2.G Kingdom/phylum-specific model training

To assess whether a kingdom/phylum-specific model can outperform the general-purpose models in any of the tools, we used the PanTEon’s module of library extraction to get a TE library by kingdoms or phyla. Then, we exclusively retained those TEs belonging to the superfamilies present in each taxonomic group with at least 10 sequences. The final dataset used had 97,462 sequences for Animalia, 63,085 sequences for Chordata, 24,126 sequences for Arthropoda, 117,493 sequences for Plantae, 114,515 for Angiosperma, 15,477 sequences for fungi, 6,312 for Ascomycota, and 5,999 for Basidiomycota. Next, using PanTEon’s Training Module, we created kingdom/phylum-specific models for each ML/DL architecture integrated in the framework (the same nine described above).

### 2.10 Model training to Distinguishing TEs from non-TE sequences

To illustrate how the PanTEon Framework can be used to generate models for any TE classification task, we chose the problem of detecting false positive sequences in TE libraries, which is a typical component of the manual curation process. To do this, we defined a binary classification problem where each model must classify a DNA sequence as either a TE (positive class) or a non-TE (negative class). To test this, we used the negative dataset generated by (66) which comprises CDS and several types of RNAs (mRNA, tRNA, rRNA, pre-mRNA, miRNA, snRNA, snoRNA, siRNA, and lncRNA) from 195 plant species. However, because that dataset was defined to be the negative class for LTR retrotransposon detection, it also contains TE elements from the other orders (such as LINE, SINE, TIR, and so on). Hence, we removed all TEs based on their sequence identifiers or BLASTn similarity to other TEs (identity > 80% and coverage >80%). The original negative dataset is available at https://zenodo.org/records/4543905. We next sought to evaluate the ability of the models to distinguish TEs from other classes of repetitive sequences. To this end, we used the raw repeat libraries generated by RM2 and available through Ensembl, from which we extracted all satellite and simple repeat sequences. This yielded 19,699 repetitive non-TE sequences. To construct a balanced negative class, we randomly sampled an equal number of CDS and RNA sequences, resulting in a total of 39,398 negative examples. For the positive class, we randomly sampled 39,398 TE sequences from the Plantae collection of the PanTEon Database, thereby generating a balanced dataset for model training and evaluation.

Finally, to assess the ability of the models to discriminate TEs from protein-coding genes, we constructed a negative control dataset using BUSCO genes identified in three plant genomes: *A. thaliana* (GCF_000001735.4), *O. sativa* (GCF_034140825.1), and *Z. mays* (GCF_902167145.1). BUSCO v.6.0.0 (67) was run in genome mode (--mode genome) using the *viridiplantae_odb12* lineage dataset. The nucleotide sequences corresponding to all complete single-copy BUSCO genes were extracted from each genome and merged into a single FASTA file containing 2,201 sequences. The PanTEon inference module was then applied to classify these sequences, and any gene predicted as a TE was considered a false positive.

### 2.11 Explainability analysis

To investigate the biological features driving TE classification, we developed a model-specific interpretability pipeline based on Integrated Gradients (IG). The analyses were performed using an in-house Python script, interpret_panteon_models.py, which is publicly available in the PanTEon GitHub repository under the bench_scripts directory. A representative subset of 160 sequences (20 randomly selected sequences from each of the eight TE orders) was extracted from the PanTEon Database and analyzed independently with the four best-performing classifiers (NeuralTE, CREATE, Terrier and DeepTE). For each model, predictions were first generated using the corresponding trained model, and IG attribution scores were subsequently computed with respect to the predicted class. For TensorFlow-based architectures (NeuralTE, CREATE and DeepTE), IG was implemented using TensorFlow’s automatic differentiation framework, whereas for the PyTorch-based Terrier model, attributions were computed using the Captum implementation of Layer Integrated Gradients over the nucleotide embedding layer. Baseline inputs consisted of zero-valued feature vectors for feature-based models and zero embeddings for sequence-based models.

Model-specific attribution scores were aggregated across all analyzed sequences to quantify the contribution of individual input features. For NeuralTE, feature importance was summarized according to the four predefined feature groups (internal *k*-mer frequencies, terminal *k*-mer frequencies, protein domain features and sequence-end features). For CREATE and DeepTE, attribution scores were computed independently for each network input branch, allowing the relative importance of *k*-mer frequency representations and one-hot encoded nucleotide sequences to be evaluated separately. For Terrier, nucleotide-level attribution scores were projected onto sequence positions through the embedding layer to identify the regions contributing most strongly to hierarchical predictions. The resulting attribution profiles were visualized as feature importance plots and summarized as average absolute attribution scores, enabling direct comparison of the biological signals exploited by the different deep learning architectures.

## 3. Results

### 3.1 PanTEon covers previous gaps in the development of transposable element classifiers

#### PanTEon Database, an extensive automatically curated dataset encompassing a large taxonomic diversity of species

The lack of a sufficiently large, publicly available and high-quality dataset of TE consensus sequences severely limits the potential for training accurate neural networks for the prediction or classification of TEs. To address this limitation, we released the PanTEon Database (release 1.6.2) composed of 239,946 TEs across 2,790 animals, plants, and fungal species (Table 1). The Database was designed to integrate curated sequences from Dfam 3.9 (386 species and other taxonomic groups) complemented by TE sequences from uncurated Dfam 3.9, APTEdb, Ensembl 2025, and B-GUT, which were predicted by RM2, automatically curated by MCHelper, and selected for being high-quality and structurally complete.

**Table 1:**
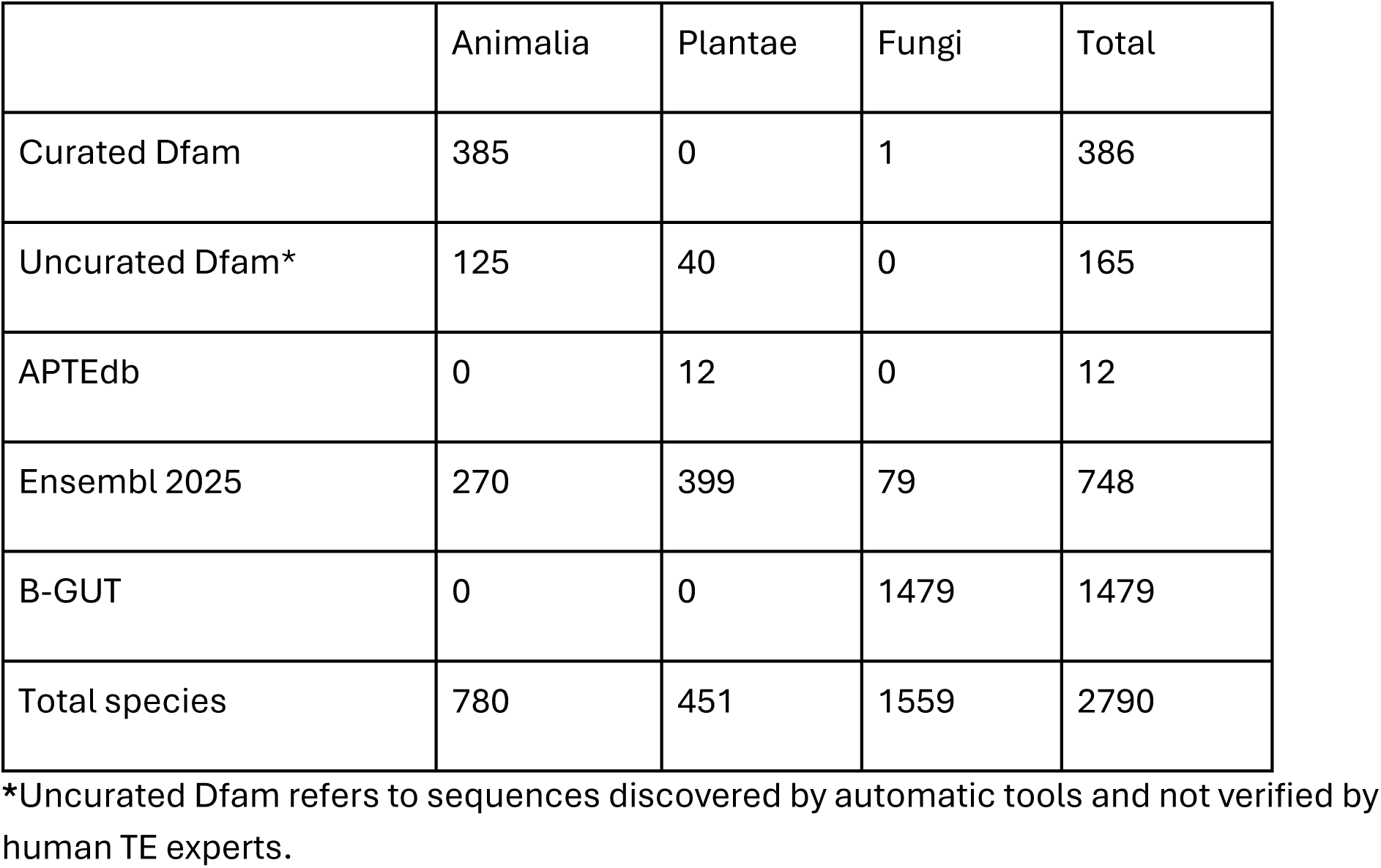
Source of species-specific TE libraries, grouped by taxonomic kingdom. A detailed list can be consulted in Supplementary File 1.

In total we added 2404 new species (from 33 phyla), corresponding to 819 new families and 332 new orders, providing the PanTEon Database with unprecedented taxonomic breadth and resolution across animals, plants and fungi (Figure 1).

**Figure 1.**
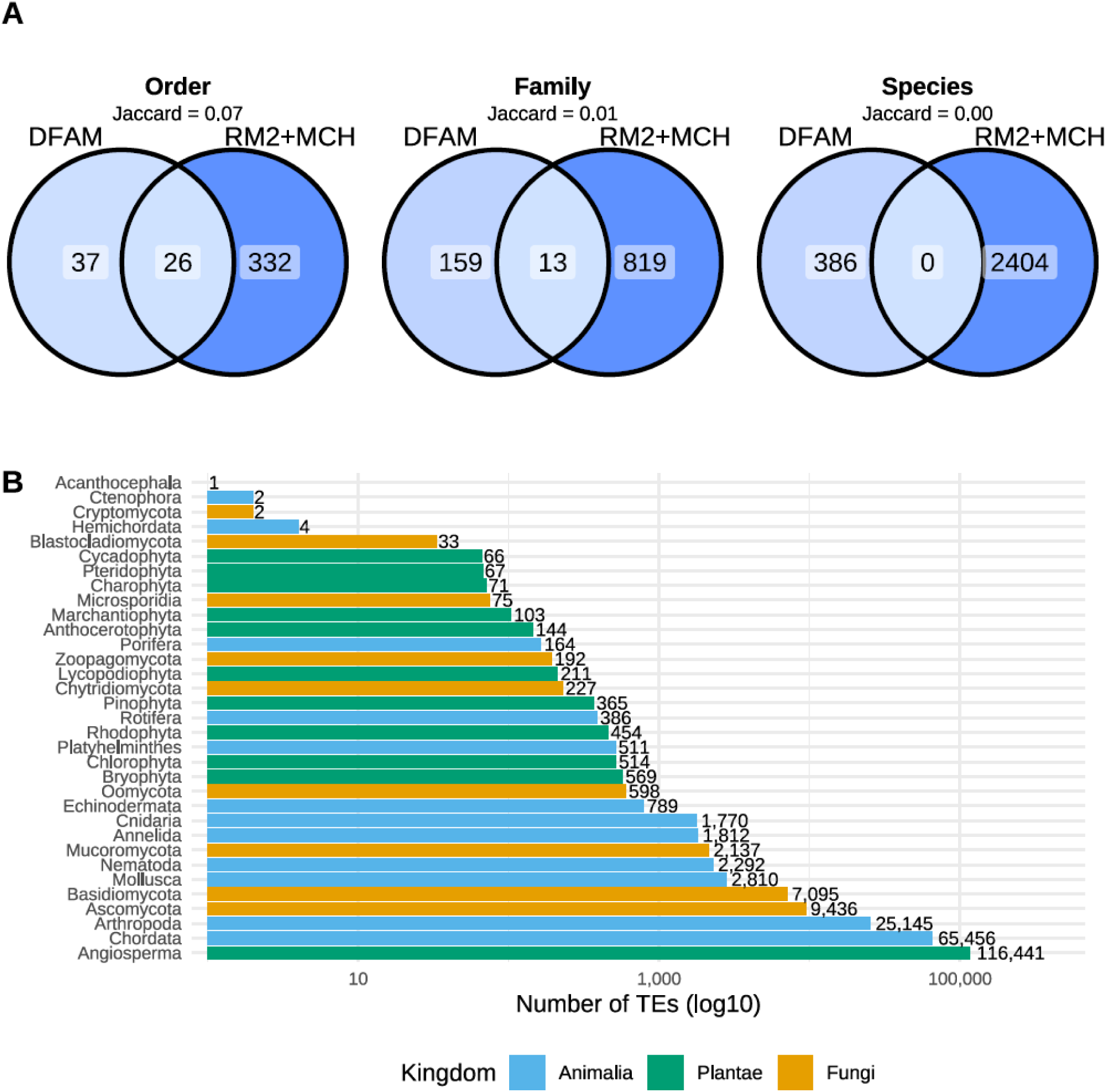
Diversity of species used to build PanTEon Database. (A) Number of unique taxonomic groups in Dfam and RM2+MCH (detected by RepeatModeler2 and automatically curated by MCHelper) sources. (B) Number of TE sequences per phylum, grouped by kingdom in the PanTEon Database.

Most of the TE sequences integrated in PanTEon Database were obtained from Ensembl 2025 with more than 160,000, followed by curated Dfam and Uncurated Dfam.

Interestingly, we found no curated sequences from plants, and only one species from fungi in Dfam (Table S6). Altogether, PanTEon Database contributes 208,354 (out of 239,946) new automatically curated TE sequences, which have evidence of structural completeness.

Although many of the TE sequences included in the PanTEon Database originate from established databases and are already available for reference-based genome annotation, this Database constitutes a valuable resource by integrating and standardizing sequences from multiple sources and applying a common curation procedure, as well as a harmonized classification nomenclature, potentially providing a more consistent reference collection across taxa. This unified collection may support the annotation of new genomes (see one example in Figure S1 and Supplementary Information), help to classify new TE sequences, especially for underrepresented species (like some fungal lineages), do large-scale comparisons across kingdoms and to be used as a training dataset to improve existing algorithms or for new ML/DL approaches to classify, curated, and annotated TEs in massive sequencing projects.

### PanTEon Database encompasses a large taxonomic diversity of TEs

Following the dynamics shown by TEs in different species, PanTEon Database is also unbalanced in terms of TE orders and superfamily quantities (Figure 2). The most abundant TE order is LTR, with 50% of all sequences, being 25% of the animal TEs, the 72% of plant TEs and 40% in fungi sequences. LINE retrotransposons are the second most frequently found TE order, being 28% of the entire dataset, but achieving up to 44% and 31% in animal and fungal sequences. In contrast, the rarest TE orders found were Maverick, Crypton, Retroposon and PLE, being only 0.024% (58 sequences), 0.047% (112 sequences), 0.047% (113 sequences), and 0.14% (344 sequences). This finding reflects the complexity of finding TEs from those orders using automatic tools, as well as their probably low natural frequency. It is important to stress that the PanTEon Database contains only structurally intact TE elements (as shown by their length distribution in Figure S2).

**Figure 2.**
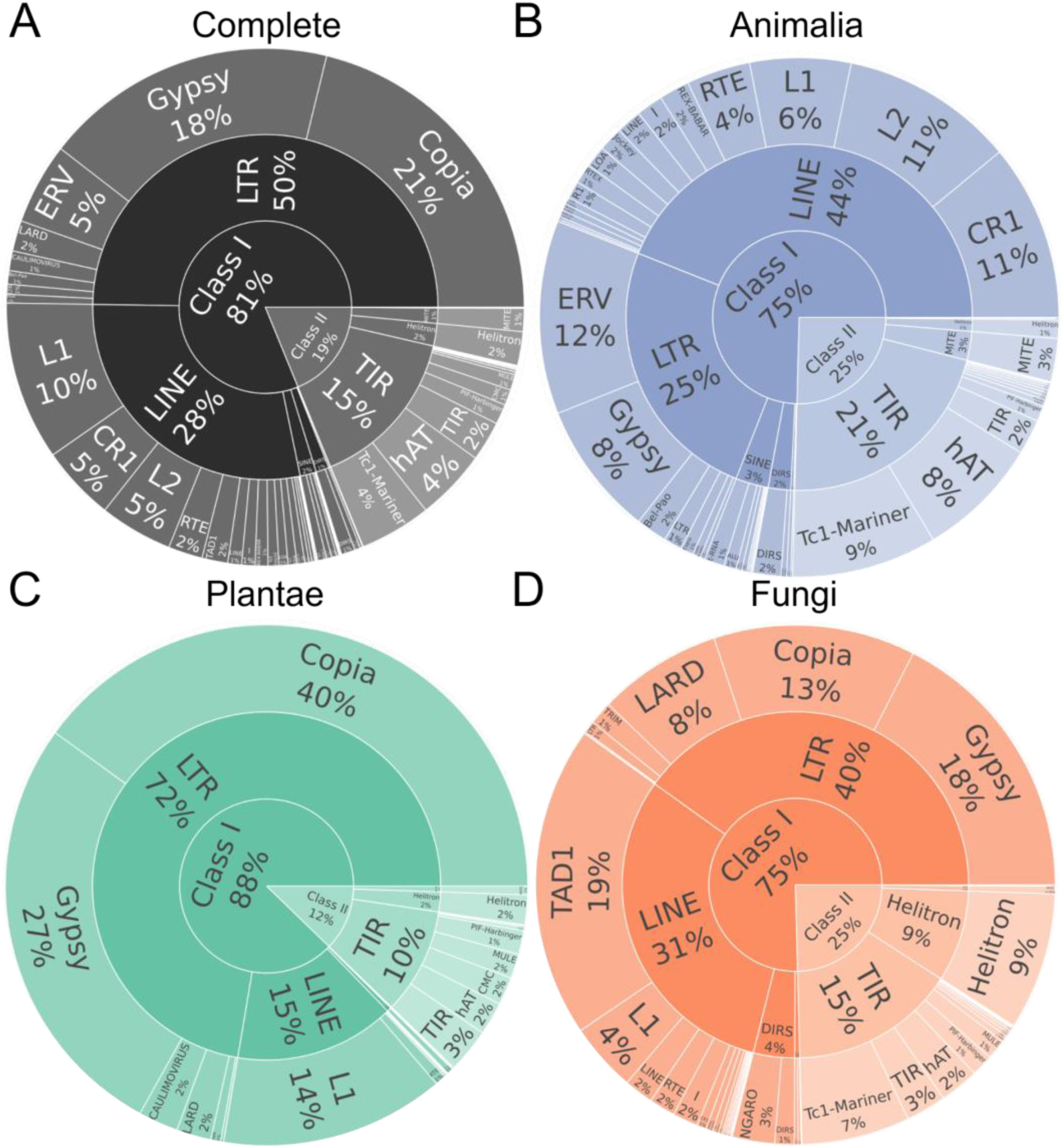
Sunburst plots showing the TE taxonomically diverse presented in the PanTEon Database, corresponding to A) all the species contained in the database, B) Animalia, C) Plantae and D) fungi.

### Overview of the PanTEon Workflow Architecture

To overcome current limitations encountered by researchers during the development, application, and evaluation of ML/DL-based tools, we released the PanTEon platform, a modular software framework that allows users to utilize state-of-the-art ML/DL algorithms to classify their TE libraries using a single tool (chosen by the user) or, alternatively, by combining several tools in parallel (Figure 3A). In the latter case, the user will obtain a combined report integrating the predictions of all selected tools. Thus, the user can compare the results and make informed decisions such as keeping only the sequences whose classification is supported by at least a certain number of tools. Moreover, if a user has a manually curated library of a species (or group of species) of interest, the user can use it to benchmark all the tools and select the best performing ones. PanTEon also allows users to create their own models based on a provided fasta file (Figure 3B). Therefore, if a user has several TE sequences and wants to specialize several models for their specific group of organisms, the user can provide them to PanTEon’s training module, which will train and create models for the algorithms chosen by the user. Thus, users can create specialized models for a specific phylum or family of species, for classifying plant LTR retrotransposon lineages, or for classifying TIR superfamilies. In fact, this module can create trained models for any classification task. The only requirement is to provide the desired categories in the sequence IDs, following the “#” character. In this way, users can create models for distinguishing between autonomous and non-autonomous elements, between TEs and non-TE sequences (as we showcase at the end of the next section), or any other potentially useful idea that a user can think of. Trained models produced by this module are compatible with PanTEon’s inference module, so users can integrate these models as new functionalities to their pipelines.

**Figure 3.**
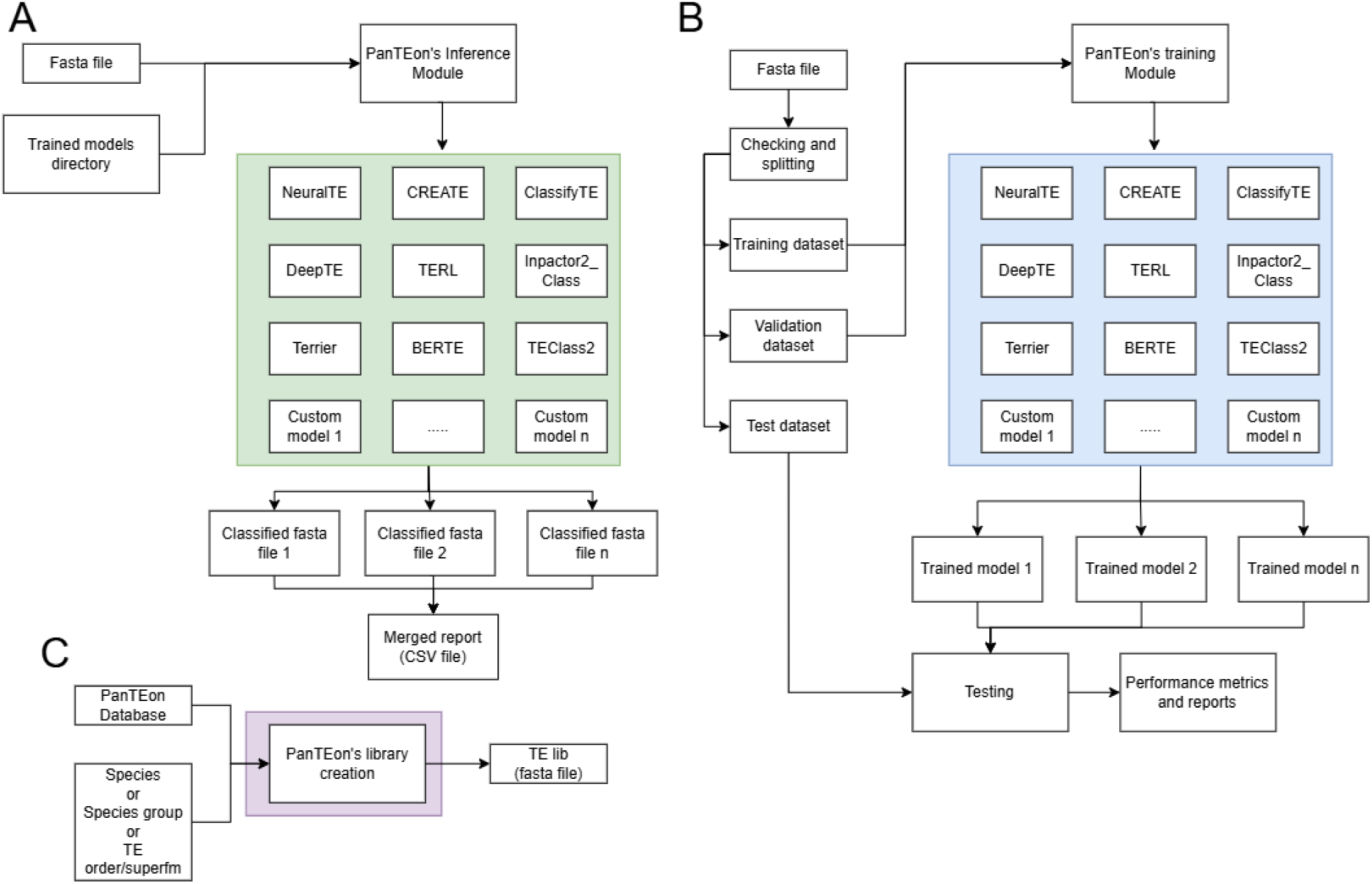
Workflow of PanTEon’s modules. A) PanTEon’s Inference Module, used for predicting the classification of TE sequences contained in a fasta file supplied by the user. B) PanTEon’s Training Module, used for retraining the models using a fasta file given by the user. This fasta file should contain the classes that will be predicted in the sequence IDs, after a “#” character. C) PanTEon’s library creation Module, used for generating TE libraries from PanTEon Database and for a specific species, taxonomic group (like kingdom, phylum, etc) or for a specific TE classification. Notice that this module offers the option to get the number of sequences for a given species, taxonomic group or TE classification in an informative way using the option --view_only.

PanTEon is also flexible, allowing users to develop their own ML/DL architectures and easily integrate them into the platform. The requirements are that the architectures must use TensorFlow or PyTorch as DL frameworks, the architecture must have three defined functions (see Figure S3 for more details), it must be developed in a Python script, and it must be placed in the “Custom_classifiers” folder. Every time that PanTEon is executed (inference or training module) the Python scripts within that folder will be imported by the framework and will be available for use inside PanTEon (for more detailed instructions on how to develop a PanTEon-compatible architecture please refer to the GitHub repository). In this way, PanTEon facilitates the development of new algorithms by allowing users to test and benchmark the new algorithms against several others in a standardized environment, with the same database, the same number of target classes, and the same metrics. Finally, the PanTEon framework may serve as a public repository of ML/DL architectures, and the developer community can contribute to PanTEon by uploading their compatible Python script, making it available to anyone in the TE community.

PanTEon can also be used for generating TE libraries. Since the computational cost of annotating a genome with a library with thousands of TEs is prohibitively high, users can use the PanTEon library creation module (Figure 3C) to extract only the required sequences. Several taxonomic groups are accepted (i.e. kingdom, phylum, class, order, family or species), as well as any TE classification: class, order or superfamily. This way, PanTEon allows users to create their own datasets from PanTEon Database to be used in masking (before annotating genes), TE annotation in closely related species, or for training specialized models.

### 3.2 The PanTEon framework facilitates the benchmarking, design and generation of DL-based model across classification tasks

#### Benchmarking based on PanTEon Database elucidates kingdom and superfamily dependency of the performance of existing classification tools

Despite growing interest in the study of TEs and in the development of AI-based TE classification tools, there is so far no independent benchmarking framework that would allow users to decide the best algorithm for their specific requirements. Thus, we used PanTEon Database as an independent dataset for benchmarking seven currently available TE classification tools: ClassifyTE, CREATE, DeepTE, NeuralTE, TEclass2, TERL and Terrier. In brief, we executed models provided by each program with default parameters, following indications in their respective GitHub repositories. We used them to classify a balanced dataset, combining all TE sequences from fungi (12,914) and identical numbers of randomly selected TE sequences from animals and plants, hence totalling 38,742 TEs, ensuring that performance comparisons were not influenced by differences in kingdom representation.

We then mapped the resulting classifications to the PanTEon Database nomenclature. All programs performed well when distinguishing Class I elements from Class II, with F1-scores ranging from 77% to 92% (Figure 4A). Classification performance started to decrease at the order and superfamily levels. At these levels, the best-performing tools were NeuralTE and Terrier, achieving F1-scores of 87% and 82% at order level, respectively, and the first obtaining 80% while the second 78% at superfamily level. This performance is somewhat lower than that reported by the authors in the tool publications, where NeuralTE achieved an F1-score of 89%, while Terrier got an average accuracy of 94%. This discrepancy shows the relevance of using independent, taxonomically diverse datasets to assess the overall performance of a tool, and that there is room for improvement even in the best-performing models, especially when dealing with species distant from those used for training these models.

**Figure 4.**
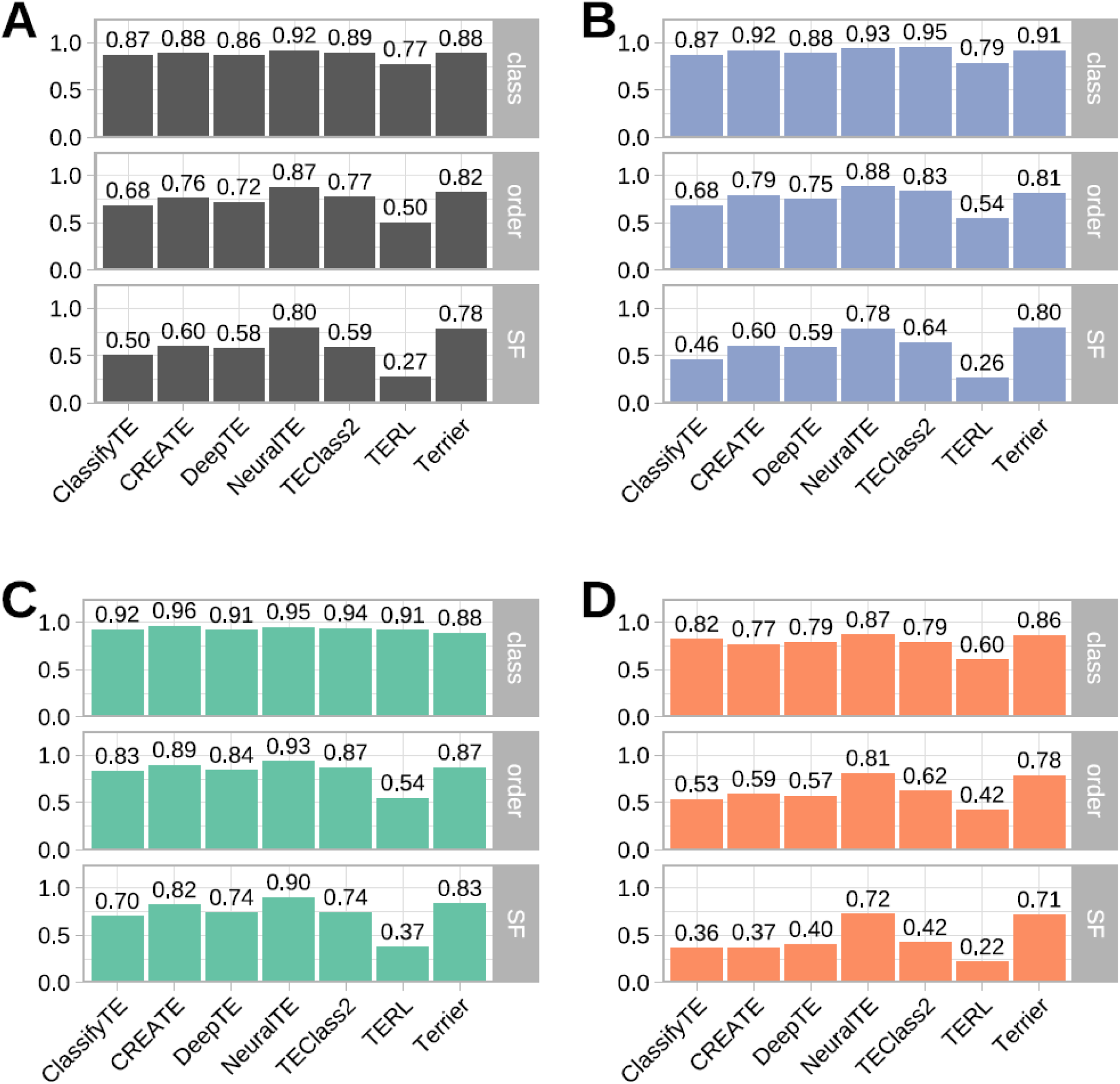
Classification tool F1-score performance for A) the entire dataset, B) animals, C) plants, and D) fungi.

We then assessed the performance of the tools for each of the kingdoms separately, to evaluate the impact of training dataset bias as most tools had been trained on Dfam and Repbase, which are heavily enriched in plants and animals. Performance for the classification at class level was higher for animals (between 79% and 95% F1-score) and plants (between 88% and 96% F1-score) than in the combined dataset (F1-score between 77% and 92%). In contrast, the performance in fungi was lower (between 60% and 87%, Figure 4B-D). A similar trend was observed at the order and superfamily levels, with most tools decreasing their performance in fungi, in some cases below 42% F1-score. Again, NeuralTE and Terrier were the two best-performing tools in the fungal dataset, with F1-scores of 72% and 71%, respectively.

Considering that each tool uses its own approach to transform DNA sequences into numerical features, as well as different ML/DL models or architectures, we were interested in their balance between performance and runtime. We found that the most common architecture type used was convolutional neural networks (4 of the 7 tools), while the ways to extract features were more variable, with *k-mer* frequency counts and DNA embeddings being the most common (5 out of 7 tools; Figure S4). Notably, the two best-performing tools (NeuralTE and Terrier) used very different approaches: while NeuralTE uses a combination of *k-mer* frequencies and structural features calculated from the TE sequences, Terrier uses a very simple approach consisting of mapping each nucleotide into an integer number (1 to 4). This allows Terrier to obtain results much faster (0.32 hours, Table S7) than almost any other tool, except for TERL which uses one-hot representation. On the other hand, although NeuralTE demonstrates a very good performance across kingdoms, the computational cost is also high (1.26 hours, Table S7).

We next focused on the specific performance of each tool across different TE superfamilies (Table S4). Because each tool predicts a different number of superfamilies, overall performance decays when the benchmarking dataset includes a superfamily (e.g. Crypton elements) that a given tool was not trained to detect. We observed that the tool with the lowest number of included superfamilies was TERL with 11 (of 28), followed by TEclass2 with 16, whereas DeepTE, NeuralTE, and Terrier included most superfamilies, with 25, 24, and 24 respectively. Although the number of considered superfamilies is directly influencing a tool’s performance in our benchmark, we consider it is crucial to use a broad benchmark dataset, which better reflects real diversity. The use of a tool not accounting for all the families putatively present in a genome is likely to result in mis-classification and low sensitivity, which will be reflected in our benchmark. Only six superfamilies were included in all the tools: LTR/Copia, LTR/Gypsy, LTR/Bel-pao, Line/L1, TIR/hAT, and TIR/TC1-Mariner. NeuralTE is the best-performing tool in the first four (all from Class I - retrotransposons), while Terrier is the best in the other two (Figure S5A). We hypothesize that the structural features used by NeuralTE allow it to better detect retrotransposons (which have terminal repeats - LTR for LTR retrotransposons and PolyA tails for LINEs - and many coding domains - up to six in LTR retrotransposons and up to 4 or 5 in LINEs). Terrier also demonstrated a very good performance when classifying sequences belonging to special supergroups, which contained sequences of a given order (LTR, LINE or TIR), but without further superfamily specification. Finally, TEclass2, which uses a Transformer-based approach, was the best-performing tool to detect one of the less common TE orders Crypton, as well as the non-autonomous SINEs. Since these orders contained the lowest number of sequences (Figure S5B), it is uncertain whether this model could generalize well in new datasets.

Overall, considering all the superfamilies, Terrier was ranked as the best tool based on the Friedman test (followed by Nemenyi post-hoc analysis), followed by NeuralTE, DeepTE, and CREATE, without significant differences between them (Figure S5C). TEclass2 and ClassifyTE showed intermediate performance, whereas TERL showed the lowest performance.

To explore the potential of integrative approaches, we combined the predictions of the seven classifiers following three ensemble methods: (i) simple majority voting, (ii) weighted majority voting, and (iii) a supervised stacking model based on XGBoost (see Methods). The XGBoost approach resulted in the best performance at all classification levels in the combined dataset, resulting in F1-scores up to 13% higher (Figure S6A), while the simple voting and weighted voting behaved similarly. This trend was consistent for the Plantae (95% F1-score; Figure S6C), Animalia (89% F1-score; Figure S6B), and fungi (80% F1-score; Figure S6D). Compared with the best-performing individual model (Figure 4), the ensemble approaches achieved notable performance gains, with increases of 7% in F1-score on the full dataset, 9% in Animalia, 5% in Plantae, and 8% in fungi. These improvements indicate that integrating the outputs of multiple classifiers through an ensemble-based ML strategy can effectively enhance predictive performance, particularly for under-represented taxa such as fungi.

### PanTEon improves benchmarking of current ML/DL architectures and allows integrating new DL-based models

Beyond comparing existing models, our benchmarking framework can be used to gain an understanding of which ML/DL architectures consistently produce better results, information that can guide the design of future tools. To allow better comparisons among models that differ in their training approach (dataset, optimization), we used PanTEon to support a standardized workflow in which every ML/DL approach was trained in the same manner. By doing this, observed differences among models should directly relate to their feature-extraction approach and their ML/DL architecture, rather than to the training approach. For this, we used PanTEon’s library extraction module to obtain a TE library per kingdom and PanTEon’s training functionality to train architectures from the seven tested tools plus two additional DL architectures: BERTE and Inpactor2_Class (Table S8). The size of the models ranged between 502k parameters (ClassifyTE) and 75M parameters (TEclass2). Most of them were evaluated originally by authors using the F1-score metric and their performance (based on the publication) ranged between 73.9% (TERL) and 89.6% (NeuralTE). A special case was Inpactor2_Class which reported an F1-score of 91% but in the original publication was trained to classify lineages of LTR retrotransposons (13 classes) of plants.

We then used the entire PanTEon Database (Supplementary File 1), and split it into 80% for the training set, 10% for the validation set, and 10% for the test set. We trained the models on the training set and reported the F1-score obtained in the test set. Most models showed a performance on the entire PanTEon Database (Table 2) similar to that reported in their respective publications, with the exception of the transformers-based architecture, BERTE, which showed a remarkable decrease in performance (-27.9%), likely due to insufficient training dataset size.

**Table 2:** F1-score results of the architecture re-training experiments using the PanTEon Database and 30 superfamilies.

| ML/DL approach | Reported F1-score (%) | Obtained F1-score (%) | Generalization test (complete sequences) | Generalization test (incomplete sequences) |
| --- | --- | --- | --- | --- |
| BERTE | 81.8* | 53.9 | 3 | 9.4 |
| ClassifyTE | 89.2* | 87.4 | 62.4 | 46.8 |
| CREATE | 84* | **88.7** | 76.6 | 71.0 |
| DeepTE | 89.2* | **88.7** | 73.6 | 66.6 |
| Inpactor2_Class | 91** | 84.7 | 74.5 | 65.4 |
| NeuralTE | 89.6 | **92.2** | 93.4 | 75.7 |
| TEclass2 | 79 | 81.5 | 38.8 | 44.0 |
| TERL | 73.9 | 73.7 | 64.2 | 39.8 |
| Terrier | N/A*** | **89.2** | 86.3 | 70.4 |

\* These architectures used HC metrics for measuring the training performance. \*\* Trained only for LTR elements in plants. \*\*\* This tool did not show the F1-score but reported an accuracy of 94%. The scores in bold correspond to the four best performances found in this study.

We next evaluated the ability of the models to classify TEs from species that were not represented in the training dataset. This property, commonly referred to as generalization, is essential for ensuring reliable performance in real-world applications. To assess generalization, we used curated TE libraries from Dfam v4.0 derived from species absent from the PanTEon database. Because the training dataset consisted predominantly of structurally complete TE sequences, we further divided the evaluation dataset into two subsets: one containing only complete elements and another containing incomplete or degraded elements. Among the four best-performing architectures, NeuralTE (+1.2% F1-score) and Terrier (-2.9% F1-score) maintained performance comparable to that observed during benchmarking when classifying structurally complete TEs. In contrast, CREATE (-12.1% F1-score) and DeepTE (-15.1% F1-score) exhibited substantial declines in performance on these previously unseen species (Table 2). Classification performance deteriorated even further for incomplete or degraded TE sequences, with F1-score reductions of 16.5% for NeuralTE, 18.8% for Terrier, 17.7% for CREATE, and 22.1% for DeepTE. These results highlight the current limitations of AI-based TE classifiers when applied to incomplete or degraded elements and to sequences from phylogenetically distant species that were not represented during training. They also emphasize the need for more diverse training datasets and models that are more robust to structural variation in TEs.

When assessing performance across 30 superfamilies in the aforementioned experiment (Figure 5A), NeuralTE achieved the best F1-scores for most superfamilies (20 out of 30), followed by DeepTE (4 out of 30), Terrier (2 out of 30), and ClassifyTE, CREATE, and BERTE (1 out of 30). Interestingly, there was not a clear correlation between the year of publication and performance (R²=0.12). This trend was particularly affected by the two transformers-based models, BERTE and TEclass2; excluding these models increased the coefficient of determination to R²=0.35, indicating that architectural differences rather than publication year primarily drive performance variation. Although this is a promising approach, their relatively low performance may indicate that the currently available data is insufficient to efficiently train models of that magnitude (Figure 5C). Nevertheless, there is a moderate increasing trend of the performance throughout the years.

**Figure 5.**
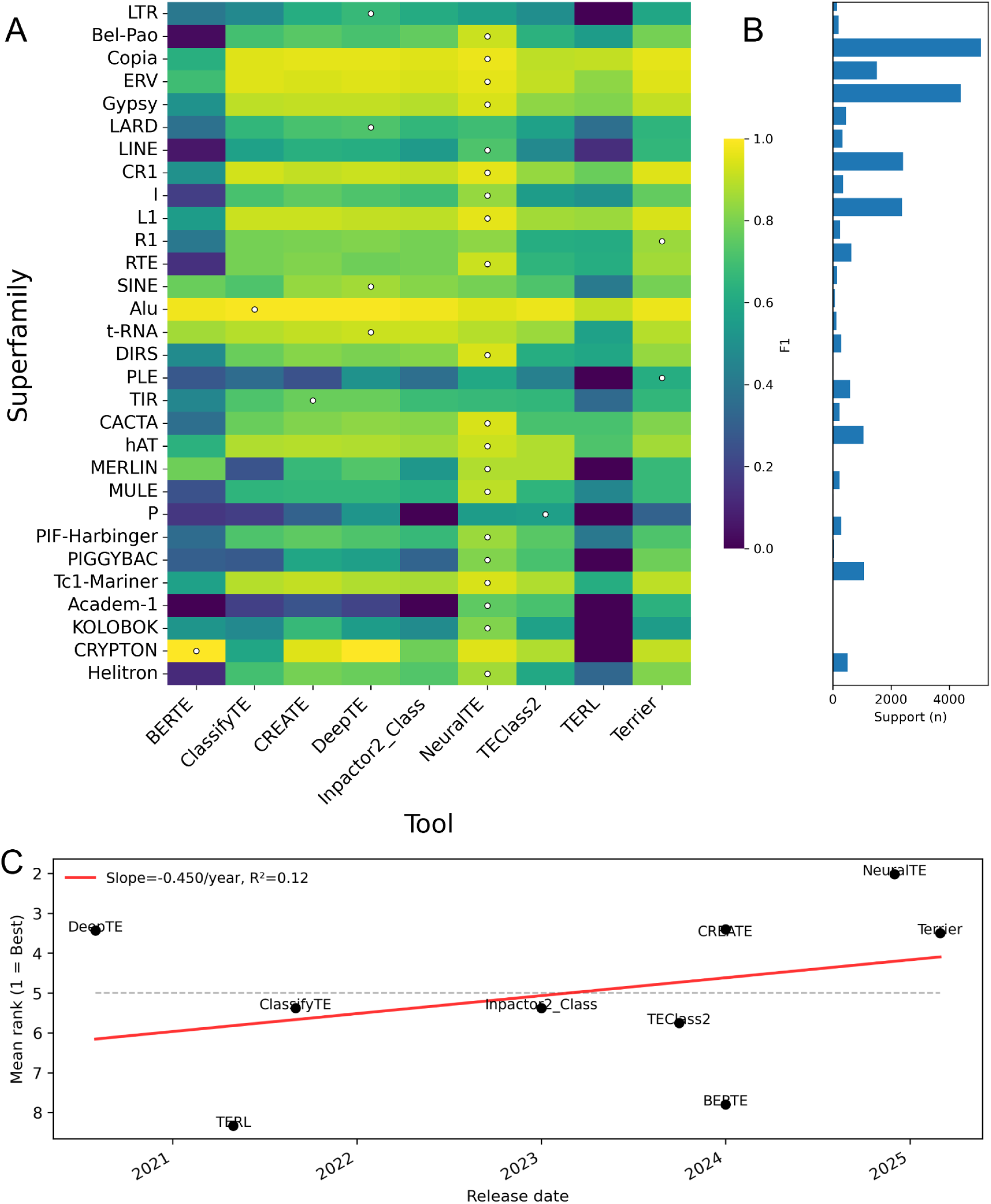
(A) F1 heatmap (rows = superfamilies sorted by TE order; columns = tools A–Z). Color encodes F1; a white dot marks the row-wise best; (B) shows the Support (n) per superfamily (number of sequences). (C) Mean rank of TE classifiers over time. Each point represents the average rank of an ML–based TE classifier across all superfamilies, plotted against its release date. Lower ranks correspond to higher overall performance. The red line indicates the linear trend (slope and R² shown in the legend). Detailed results can be consulted in Supplementary Table S9.

Finally, we assessed whether model size and computational time impacted the model’s performance. Unexpectedly, the models with better performance were not the largest ones: NeuralTE and Terrier obtained good performances with fewer than 2 million trainable parameters, whereas TEclass2 (with more than 75M) had one of the lowest F1-scores (Figure S7A). Some large models, like CREATE (>72M) did obtain a good performance.

Interestingly, ClassifyTE, one of the smallest models (502K parameters) got a reasonably good F1-score, but required more time than most of the other models (except DeepTE, NeuralTE, and TEclass2), demonstrating a huge impact of the pre-processing and feature generation steps. Actually, this model and NeuralTE were the two models that proportionally invested more time in feature generation as compared to training (Figure S7B; Table S10). This underscores the necessity to have an appropriate balance between feature generation and model scale to ensure good results in efficient runtimes, especially when analyzing large datasets.

### Generation of phylum-specific models with PanTEon’s training module

Based on the above results, we hypothesized that building a general-purpose, cross-kingdom model to predict classifications in all species is probably not the best option, since superfamilies are unevenly distributed across taxonomy. For example, it is well-known that LTR/Bel-Pao is commonly found in insects, but not in plants, and we found LINE/L1 has many sequences in the PanTEon Database for animals (6,353), but they were rarely present in fungi (849). Thus, we used PanTEon training functionality to generate phylum-specific models for all kingdoms and phyla with more than 7,000 TE sequences: Animalia, Plantae, and fungi, as well as Chordata, Arthropoda, Angiosperms, Ascomycota, and Basidiomycota. For each model, we only considered superfamilies with at least 10 sequences in the corresponding kingdom or phylum, resulting in 30 superfamilies for Animalia, 29 for Chordata, 25 for Arthropoda, 24 for Plantae, 21 for Angiosperms, 24 for fungi, 21 for Ascomycota, and 21 for Basidiomycota (Table S11).

We benchmarked the performance of these models and compared it to the general model. Our results showed different results depending on the dataset. For instance, we observed that the overall performance of all the models (except for TEclass2) was at least as high when training only with animals compared to using the general dataset (Figure 6). Similarly, when using only chordate data, six models achieved F1-score above 90%, showing an overall improvement. However, when using arthropod data, the performance decayed slightly as compared to the general model. Using data from Plantae, we noticed an increase in F1-score with respect to the general dataset, with performance increasing further when using only angiosperms. For fungi, the overall performance decayed for the specific dataset compared to the general model. Nevertheless, the overall model performance increased when using data for Ascomycota and Basidiomycota, but remained below that of the general model, probably because of the small number of fungal TE sequences contained in the Database.

**Figure 6.**
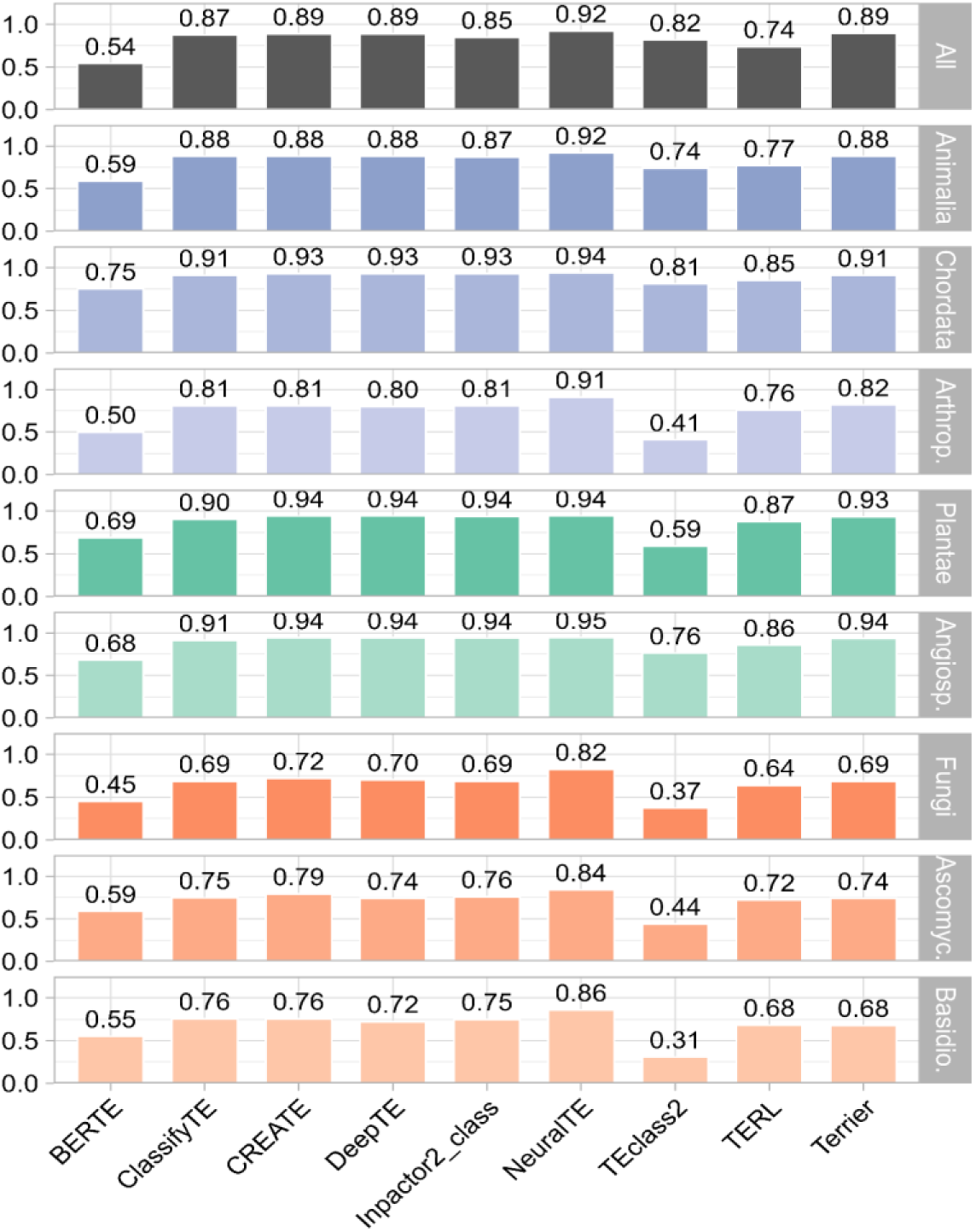
Classification performance of TE classifiers across major taxonomic groups. Bars represent the weighted mean F1-score obtained by each model within each group, with colors indicating the corresponding taxonomic category (Animalia, Plantae, fungi, etc.). Panels are organized by groups to highlight differences in classifier performance and cross-kingdom generalization ability. The detailed performance by each superfamily can be consulted in Tables S12-S19.

When we computed weighted F1-scores based on the relative proportion of TE sequences across Animalia, Plantae, and fungi, we observed notable shifts in model performance compared with the general (combined) evaluation. Transformer-based models showed clear improvements, with BERTE increasing from 0.54 to 0.63 and TEclass2 from 0.64 to 0.82. In contrast, other approaches exhibited slight performance declines, including Inpactor2_Class (−0.04) and TERL (−0.07), while the remaining models remained largely stable.

### TE versus non-TE sequence classification models across several current ML/DL architectures

After testing several aspects of the superfamily classification task in state-of-the-art ML/DL models, we now focused on using the PanTEon Framework in another classification problem. One of the most time-demanding tasks in the TE workflow is the detection and elimination of false-positive sequences, along with the overall improvement of library quality for TEs generated by automatic tools. This process, called manual curation, is performed by TE experts following several step-wise protocols, but in this case, we were interested only in filtering out sequences that do not correspond to TEs (i.e. genes, CDS, RNAs) and we asked if the models used to classify TEs into superfamilies also can be trained to do this automatically. So, we used the PanTEon training functionality again but this time using a dataset corresponding to two classes, TEs and non-TEs. For doing that, we used a modified version of the negative dataset from 195 plant species (non-TE sequences) published in a previous study (Orozco-Arias, Candamil-Cortés, Jaimes, et al., 2021), and a set of satellite repeats and simple repeats extracted from the Ensembl’s raw TE libraries (created by RM2). For the positive dataset, we used the plant TE sequences available in the PanTEon Database. For computational efficiency, a subset of 39,398 sequences was randomly sampled for each class (positive and negative).

All methods showed high predictive performance, with most of the models achieving more than 0.95 F1-score and four of them surpassing 0.97 (Figure 7A). As a negative control, we extracted the BUSCO single-copy genes found in three plant species (*A. thaliana, O. sativa* and *Z. mays*) and then we used the trained models to infer the classification between negative and positive classes. Most of the models (7 of 9) showed false-positive rates between 0 and 1%, while BERTE got 4% and NeuralTE got 6% (Figure 7B). These results are in line with the ones reported in (34), where TERL, DeepTE, Inpactor2 and CREATE achieved performance above 90% in F1-score, and also following the patterns shown in (66), where the researchers reported an F1-score up to 97.9% F1-score using anMLP algorithm.

**Figure 7.**
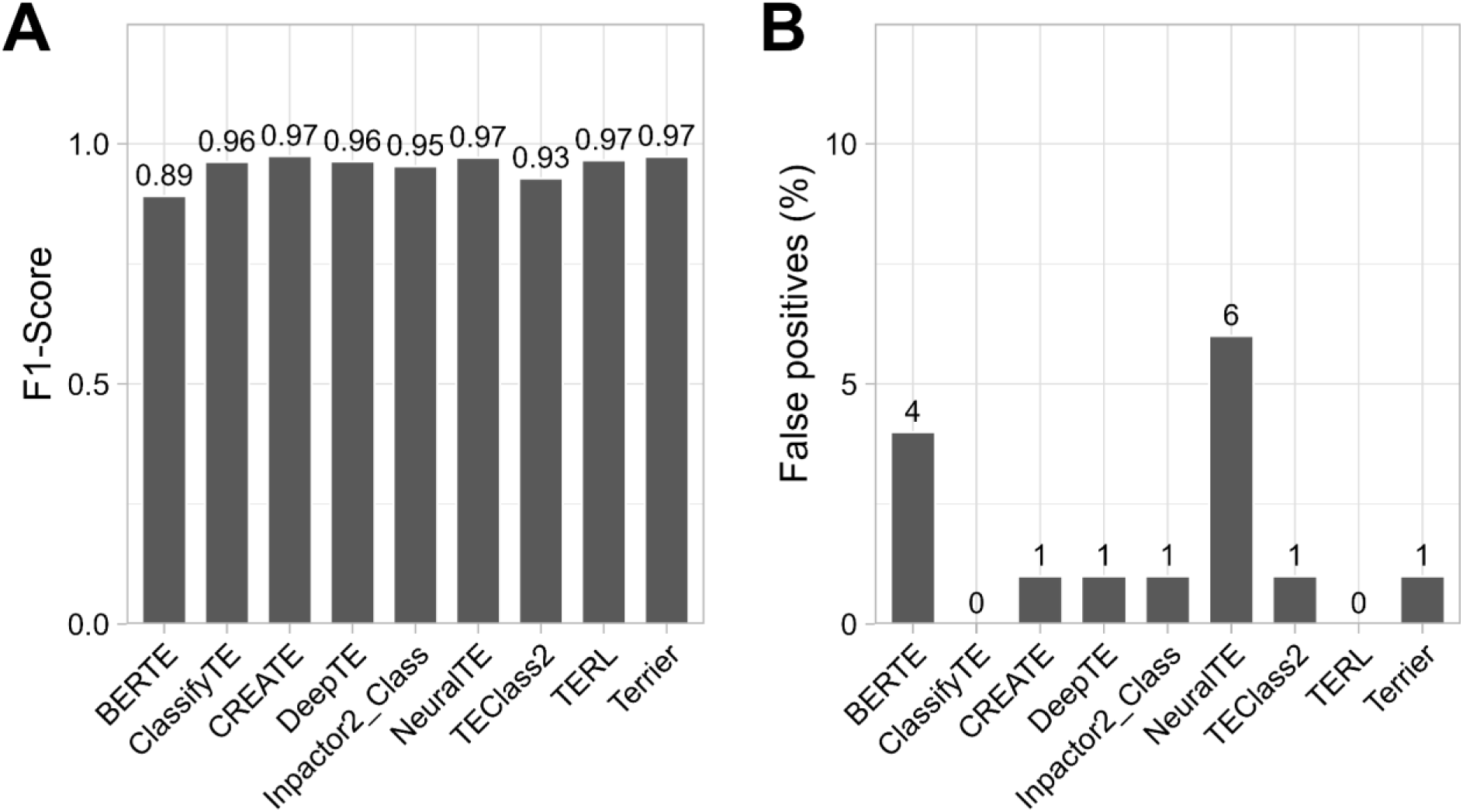
Performance of TE classifiers in distinguishing TEs from non-TE sequences. A) F1-score and B) Percentage of false-positive predictions using a negative control dataset.

Nevertheless, the second study only used LTR retrotransposons as a positive class, while in this case we used all the TE orders found in plants. Thus, using the PanTEon framework, it was possible to solve the problem of distinguishing false positive sequences in plant genomes, and the resulting trained models can be used within the PanTEon framework, through the Inference Module.

## Discussion

Accurate TE annotation remains a major challenge in modern genomics, owing to the extraordinary diversity of TE lineages and the persistent lack of standardization of detection, classification, and curation procedures. PanTEon addresses these limitations by integrating a taxonomically broad and methodologically consistent TE dataset with a unified framework for model training, benchmarking, and application. This publicly available resource, comprising almost 240,000 curated TE sequences from over 2700 species across Animalia, Plantae, and fungi, enables rigorous evaluation of existing tools and supports the development of new ML/DL models under reproducible and standardized conditions. By providing a methodologically uniform training dataset, standardized labels, and a flexible architecture for integrating custom-made classifiers, PanTEon establishes the first DL-based cross-kingdom computational ecosystem dedicated to TE research.

Our benchmarking analyses demonstrate that TE classification performance is strongly conditioned by taxonomy. Existing tools often generalize poorly across kingdoms and perform particularly badly in fungi, reflecting the limited representation, fragmentation, and annotation quality characteristic of fungal TE datasets (70). Retraining ML/DL architectures within PanTEon markedly improved performance—particularly for Chordata, Plantae, angiosperm, and Basidiomycota—showing that taxon-informed model design can significantly enhance classification accuracy. Performance differences across TE superfamilies further revealed the importance of balanced datasets, with well-represented groups such as LTR/Copia, LTR/Gypsy, LINE/L1, LINE/RTE, and TIR/TC1-Mariner consistently achieving higher F1-scores than sparsely sampled orders such as PLE, Crypton, and Maverick. These results highlight ongoing needs in TE database expansion and a potential for synthetic data generation. Although informative, our benchmarking study is limited by the lack of more independent curated datasets. Future benchmarking efforts would benefit from community-developed gold-standard datasets specifically designed for TE classification, analogous to benchmark resources available in other ML domains.

Comparing different models within a standardized framework revealed that simple sequence-derived features, such as one-hot encodings and *k*-mer frequencies, provide fast and effective representations but may fail to capture the high nucleotide-level heterogeneity of TEs, while genomic language model embeddings showed limited performance, likely due to the current scarcity of training data. In contrast, combining sequence-derived features with biologically meaningful structural characteristics, such as terminal repeats and coding-domain similarities, substantially improved classification accuracy, albeit at the cost of increased computational time, with feature extraction accounting for 98.3% of NeuralTE’s runtime (see Figure S7B and Supplementary Information).

Our interpretability analyses revealed that, despite relying on distinct feature representations and architectures, the best-performing classifiers converged on remarkably similar biological signals (see Figures S8-S9, Tables S20-S21, and Supplementary Information). Across all models, internal sequence composition emerged as the primary determinant of TE classification, whereas terminal sequence features contributed substantially only for TE orders characterized by conserved terminal structures, particularly LTR retrotransposons, TIR elements and, to a lesser extent, DIRS and Helitrons. Moreover, attribution analyses demonstrated that the models learned biologically meaningful features, including TE order-specific coding domains and lineage-specific sequence signatures, rather than relying on generic sequence patterns, although a small subset of features consistently accounted for the highest attribution scores, in agreement with previous explainable AI studies on LTR retrotransposon classification (66, 71). In particular, *k*-mer-based representations consistently provided greater discriminatory power than terminal sequence encodings, highlighting the importance of global sequence composition over local terminal motifs for distinguishing TE orders.

Collectively, these findings indicate that modern AI-based classifiers do not operate as black boxes but instead exploit evolutionary and structural characteristics that are well established in TE biology, providing confidence that their predictions are supported by biologically interpretable features rather than spurious correlations.

Convolutional neural networks consistently outperformed transformer-based architectures, suggesting that TE classification primarily depends on local sequence patterns, structural signatures, and coding domains rather than long-range sequence dependencies (see Supplementary Information). Transformer performance may improve as larger curated TE datasets and TE-specific foundation models become available.

Interestingly, model size correlated with computational time but not with classification performance, indicating that biological inductive bias and feature representation are more important than model complexity. Among the evaluated methods, Terrier provided the best balance between accuracy and efficiency through its lightweight architecture and hierarchical loss function, whereas CREATE relied exclusively on generic sequence representations, making it potentially more transferable to other genomic classification tasks. More broadly, PanTEon demonstrates that standardized benchmarking combined with explainable AI provides a robust framework for evaluating TE classifiers while revealing the biologically meaningful features underlying their predictions.

The broader implications of PanTEon extend beyond classification accuracy. By revealing the taxonomic and structural signals learned by DL models, PanTEon exposes fundamental limitations in current TE ontologies and emphasizes the need for harmonized, machine-readable nomenclature. Its unified benchmarking environment provides a scalable and evolutionarily informed basis for TE annotation in large international genome consortia, where inconsistency in TE classification remains a major bottleneck for comparative genomics. At the same time, the biologically meaningful latent representations learned by PanTEon models open new avenues for evolutionary inference, including the discovery of novel TE families, reconstruction of deep homologies, and refined detection of horizontal transfer.

The current version of the PanTEon framework is dedicated to TE classification. Nevertheless, its modular design allows the future incorporation of additional TE-related tasks, including identification (i.e. detecting TEs *de novo* inside a genome; via the --task identification parameter) and trimming of a chimeric or over-extended TE sequence (a crucial step in the curation process; via the --task trimming parameter). Although no ML/DL approaches currently address these two tasks, we anticipate that future developments will incorporate AI-based solutions. PanTEon has been designed to readily integrate such advances, as well as additional TE-related tasks.

Despite its advances, PanTEon’s performance ultimately depends on the quality of existing genomic resources, which remain uneven in taxonomic breadth and assembly completeness. Also, the PanTEon Database is unbalanced in terms of both the number of species in each taxonomic group and the number of sequences for each TE order. This imbalance primarily reflects two factors. First, some species groups have received much more attention in genomic research than others for many different reasons, and second, some species groups have much more TE diversity in their genomes (like angiosperms; Figure 1B). Another limitation of the current PanTEon Database is that it contains only structurally complete TE sequences that exhibit the expected features of their assigned classification. While this strategy improves annotation confidence and reduces label noise during model training, it does not fully capture the diversity of TE-derived sequences found in real genomes, where fragmented, truncated, nested, and highly degenerated copies are common. Consequently, models trained exclusively on PanTEon Database sequences may show reduced performance when classifying heavily degraded genomic TE fragments (Table 2). Future efforts are required to palatinally find better non-autonomous/structurally incomplete TE representatives and to include them in future versions of PanTEon Database. Nevertheless, the framework creates new opportunities for next-generation TE classification by enabling modular model integration, multimodal architectures, and the potential development of TE-specific foundation models. By delivering the first deep-learning-based framework for cross-kingdom TE classification, PanTEon provides a reproducible foundation upon which future methods can be reliably compared and sets the stage for transforming TE annotation into a mature, quantitative, and fully standardized domain within computational genomics.

## 4. Conclusion

TE annotation is still an immature area within the field of genomics. In contrast to gene annotation, the annotation of TEs still requires intensive manual work, limiting their study on a large-scale and resulting in severely limited and strongly biased knowledge. AI-based algorithms have shown promising results, especially when used to classify TEs into classes, orders or superfamilies, but current ML/DL models are restricted to only a few tasks, and there is a lack of standardized frameworks to compare their performance. Here, we present PanTEon, a framework composed of a freely accessible database of almost 240 thousand automatically curated TEs from over 2700 species (covering animals, plants and fungi) and a platform that integrates nine state-of-the-art classification algorithms.

Using the PanTEon Database, we benchmarked several TE classification tools in a fair way, obtaining relevant insights into design features associated with high performance or computing efficiency. Moreover, we demonstrated a strong taxonomical bias, particularly affecting fungal genomes. Finally, we demonstrated how, using a single command line,

PanTEon can train several ML/DL models (plus custom user-developed models) either with a general purpose, or with taxon-specific or TE family-specific focus, and tailored to TE classification or other TE-related tasks. Altogether, PanTEon establishes a foundational tool for developing, testing, and integrating ML/DL models in any TE-related task. We expect this resource to sustain a community-driven effort to transform TE annotation into a mature, quantitative, and fully comparable domain within computational genomics.

## 5. Acknowledgements

We thank the HPC bioinformatics platforms from French Bioinformatics Institute (https://www.france-bioinformatique.fr) for their HPC support. We also acknowledge the computational resources provided by the Bioinformatics and Pattern Recognition group server (BIOINFO-CP) and the multi-user cluster at UTFPR in Cornélio Procópio, Brazil.

## 7. Data availability

The PanTEon Database and the trained models to reproduce the results in this study are deposited at Zenodo. The data and models can be downloaded from Zenodo at https://doi.org/10.5281/zenodo.18039746. This includes the complete PanTEon Database version, the metadata as well as the version used for benchmarking. The software of the PanTEon platform is freely available on GitHub at https://github.com/simonorozcoarias/PanTEon.

