## Supplementary Information for "PanTEon: a cross-kingdom framework to guide the design of transposable element classifiers"

**Supplementary Figures**

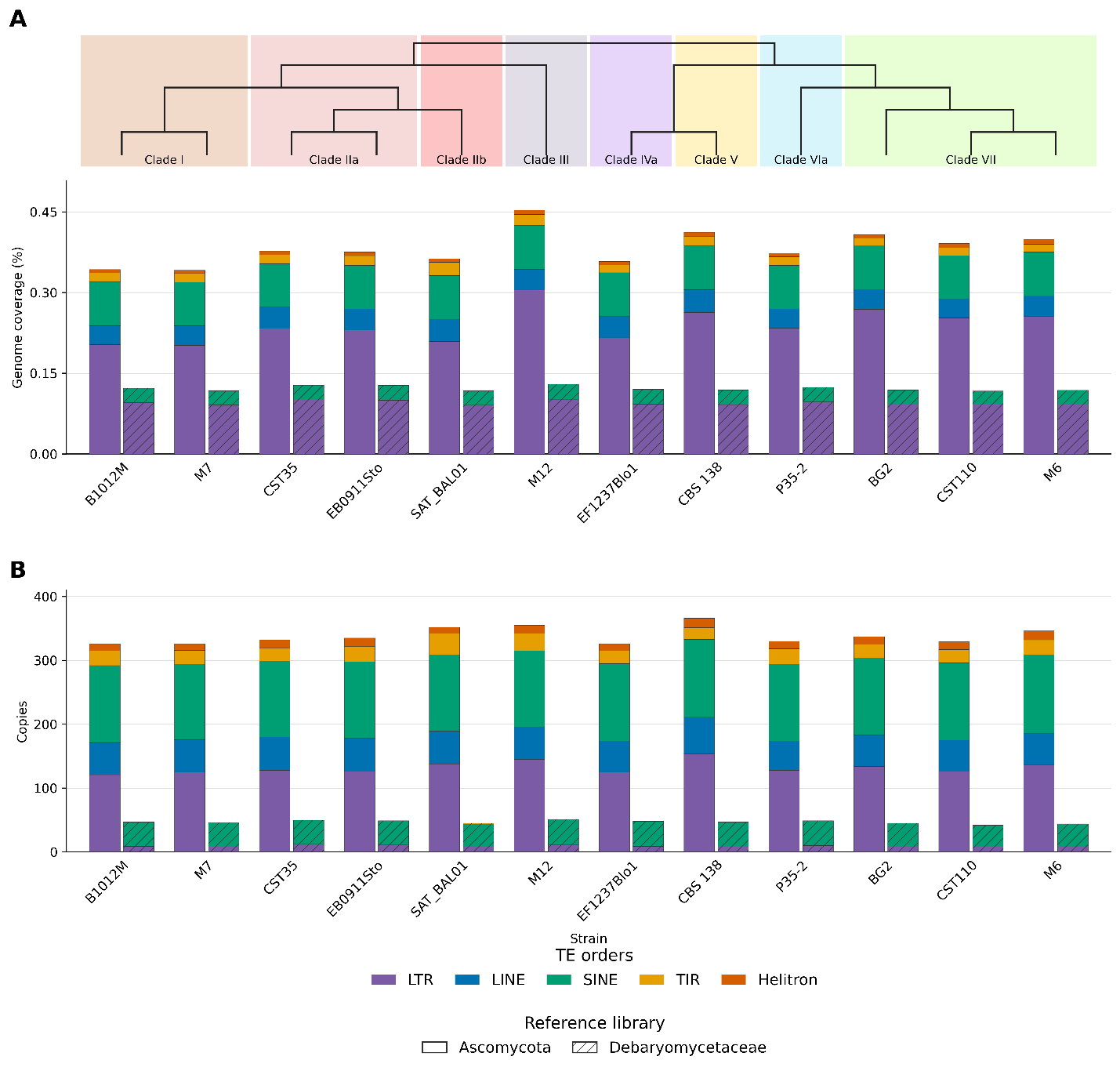

**Figure S1**. Application of PanTEon reference libraries to TE annotation in *Candida glabrata*. RepeatMasker annotations were generated using two libraries containing sequences from Ascomycota or Debaryomycetaceae extracted from the PanTEon Database. (A) Percentage of each strain’s genome covered by TE annotations, partitioned by TE order. (B) Number of inferred TE copies per strain and order; annotation fragments sharing the same RepeatMasker ID within a contig were counted once. For each strain, the left, unhatched bar represents the Ascomycota library, and the right, hatched bar represents the Debaryomycetaceae library. Colors indicate TE orders. The dendrogram above panel A reproduces the reference topology for the analyzed strains taken from (Marcet-Houben et al., 2022), with shaded backgrounds indicating clades. Branch lengths are not to scale.

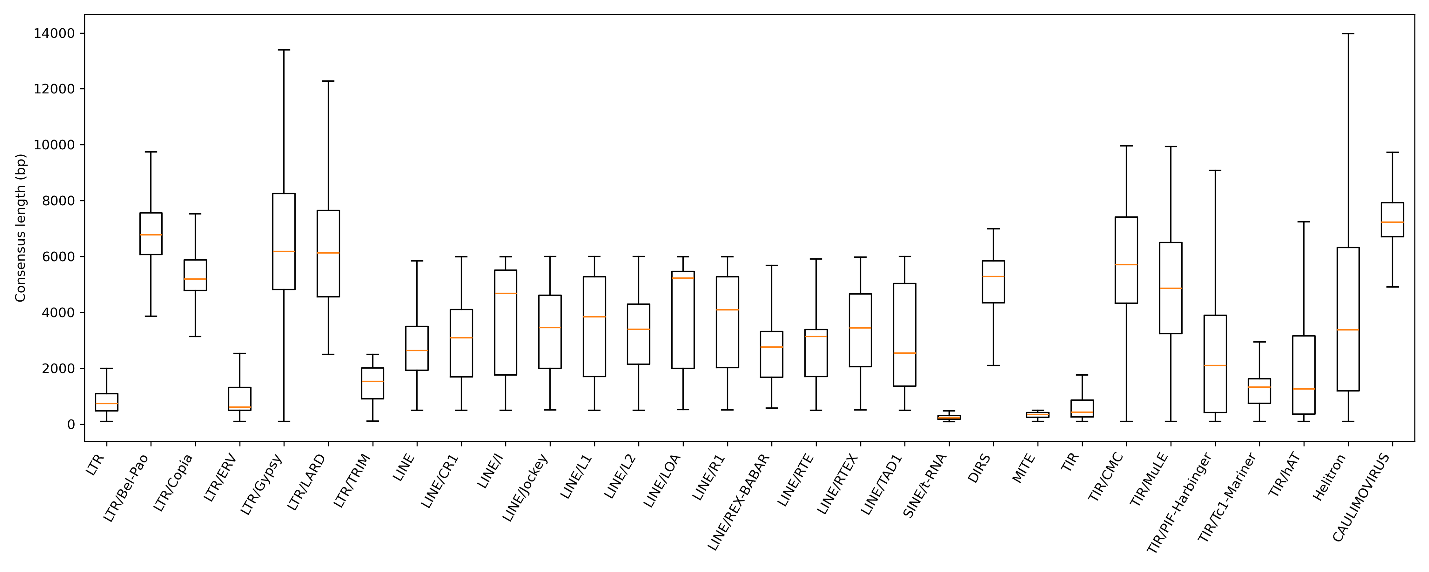

**Figure S2**. Length distribution of transposable elements across superfamilies in the PanTEon database (release 1.6.2).

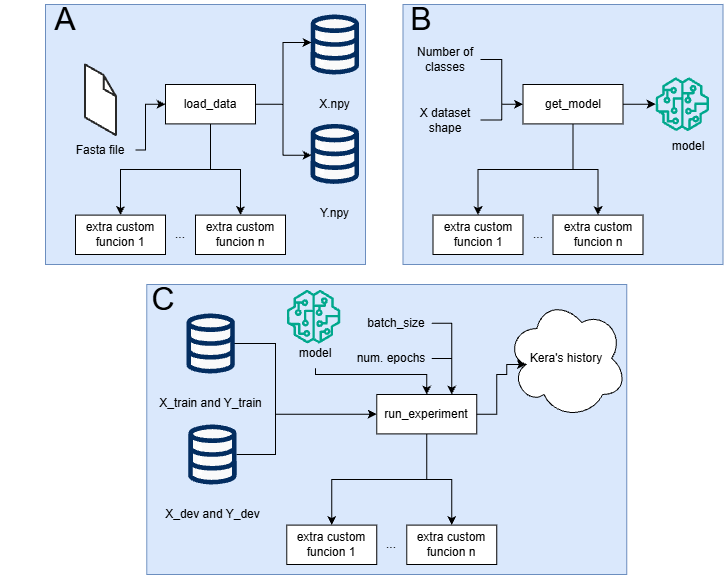

**Figure S3**. Standardized functions contained in training script used in PanTEon’s compatible architecture. A) Load data function, needed to get the data from a fasta file and extract and generate the features used for training. It should return and numpy array containing the features (X.npy in the figure) and another with the labels (Y.npy in the figure). B) Get model function: this function oversees generating the ML/DL model using whether Tensorflow, Pytorch or even Scikit-learn for traditional ML algorithms. C) Run experiment function: this function oversees training the model generated by the get_model function using the datasets generated by load_data function. Every architecture developed by third-party users should contain those three functions to be compatible with PanTEon’s modules. For more details in how to include custom architectures in the PanTEon Framework, please refer to the GitHub repository.

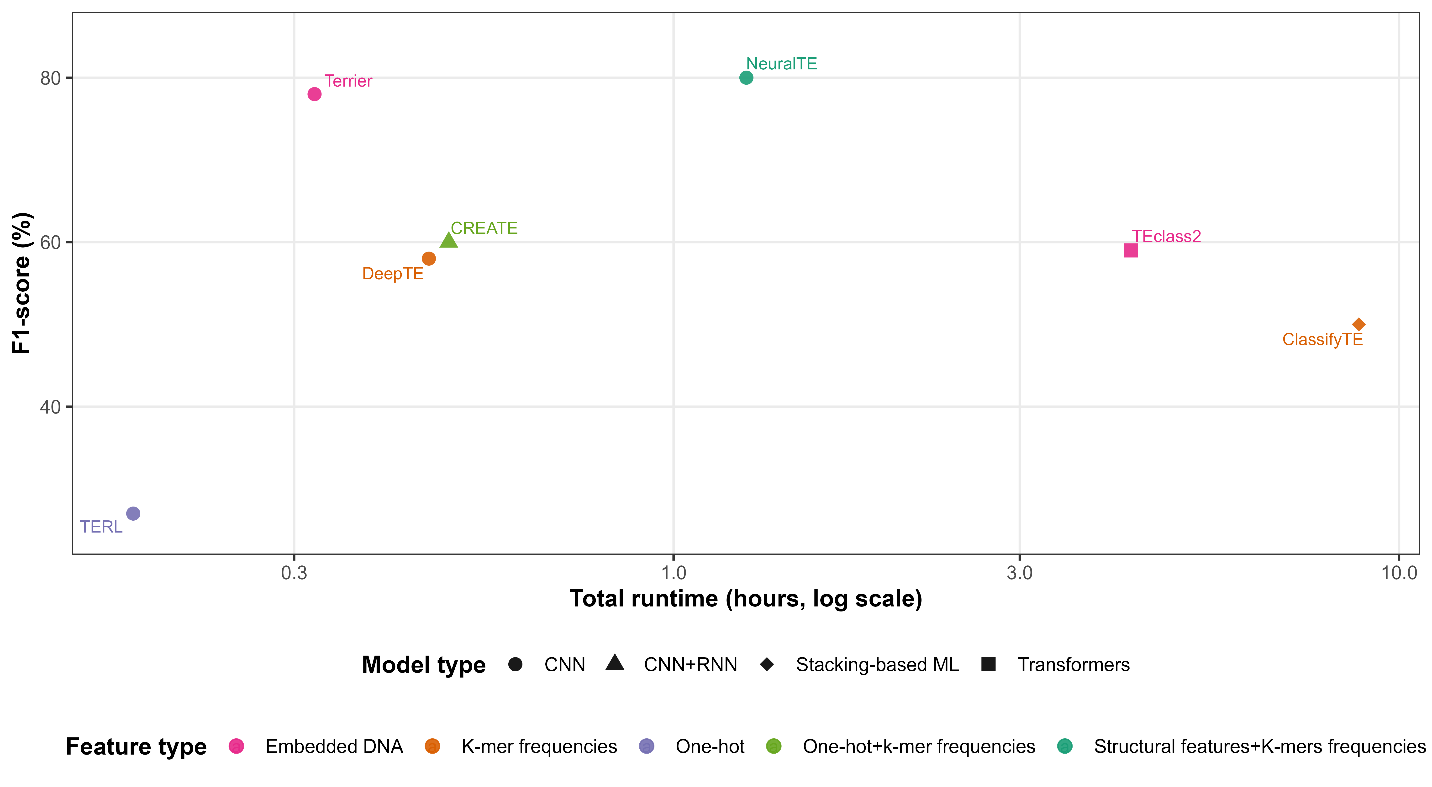

**Figure S4.** Relationship between feature type, runtime, and classification performance. Each point represents a TE classifier (shape = model type; color = feature type). The x-axis shows total runtime (log scale) and the y-axis achieved F1-score. Models using hybrid feature representations tend to achieve higher F1-scores but require longer runtimes, whereas one-hot, *k-mer* and embedded features offer more computationally efficient trade-offs. Tool Runtimes can be consulted at Table S7.

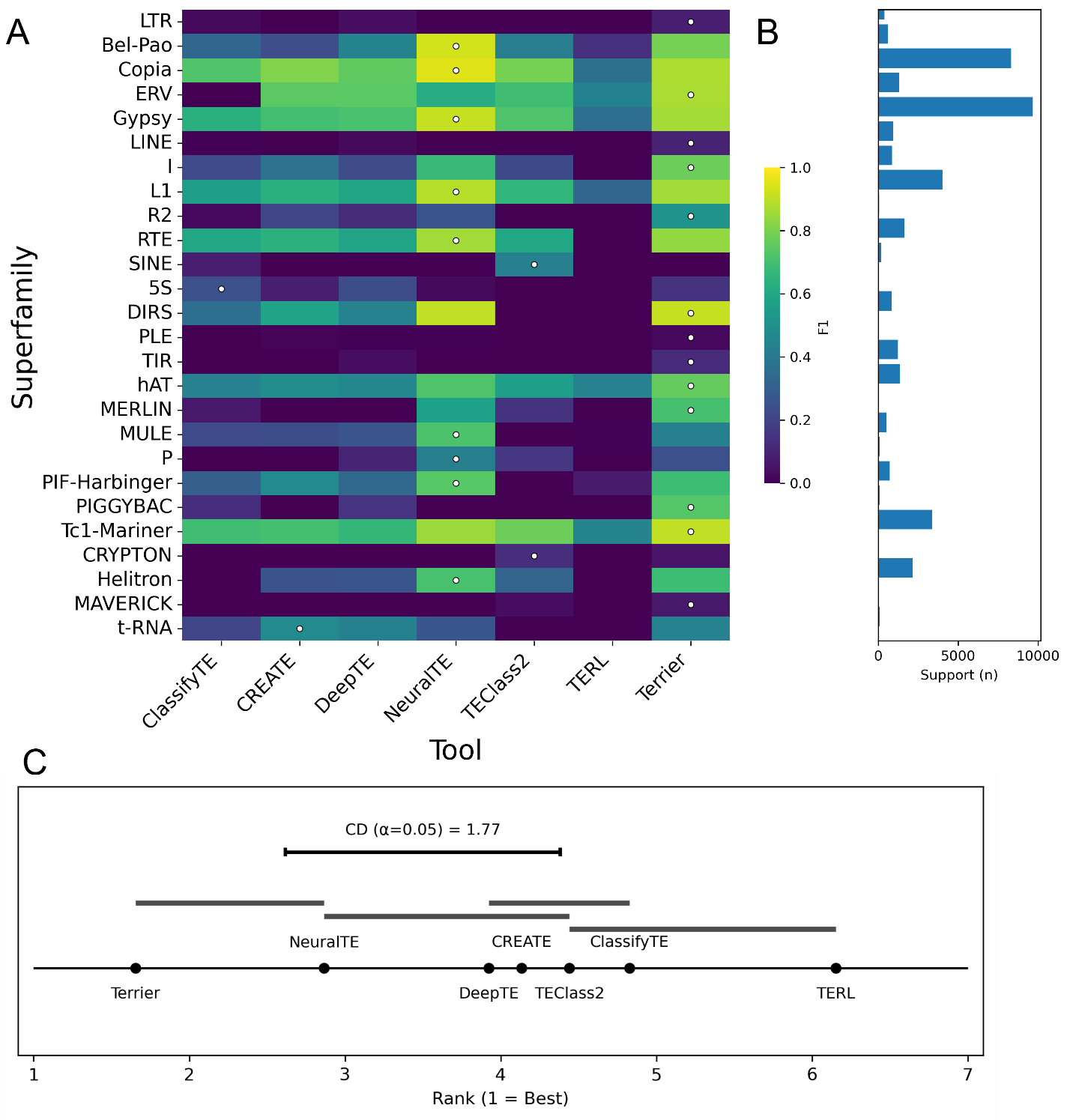

**Figure S5.** (A) F1 heatmap (rows = superfamilies sorted by TE order; columns = tools A–Z). Color encodes F1; a white dot marks the row-wise best; (B) shows the Support (n) per superfamily. (C) Critical difference diagram of average tool ranks across all superfamilies. Lower ranks correspond to higher overall performance.

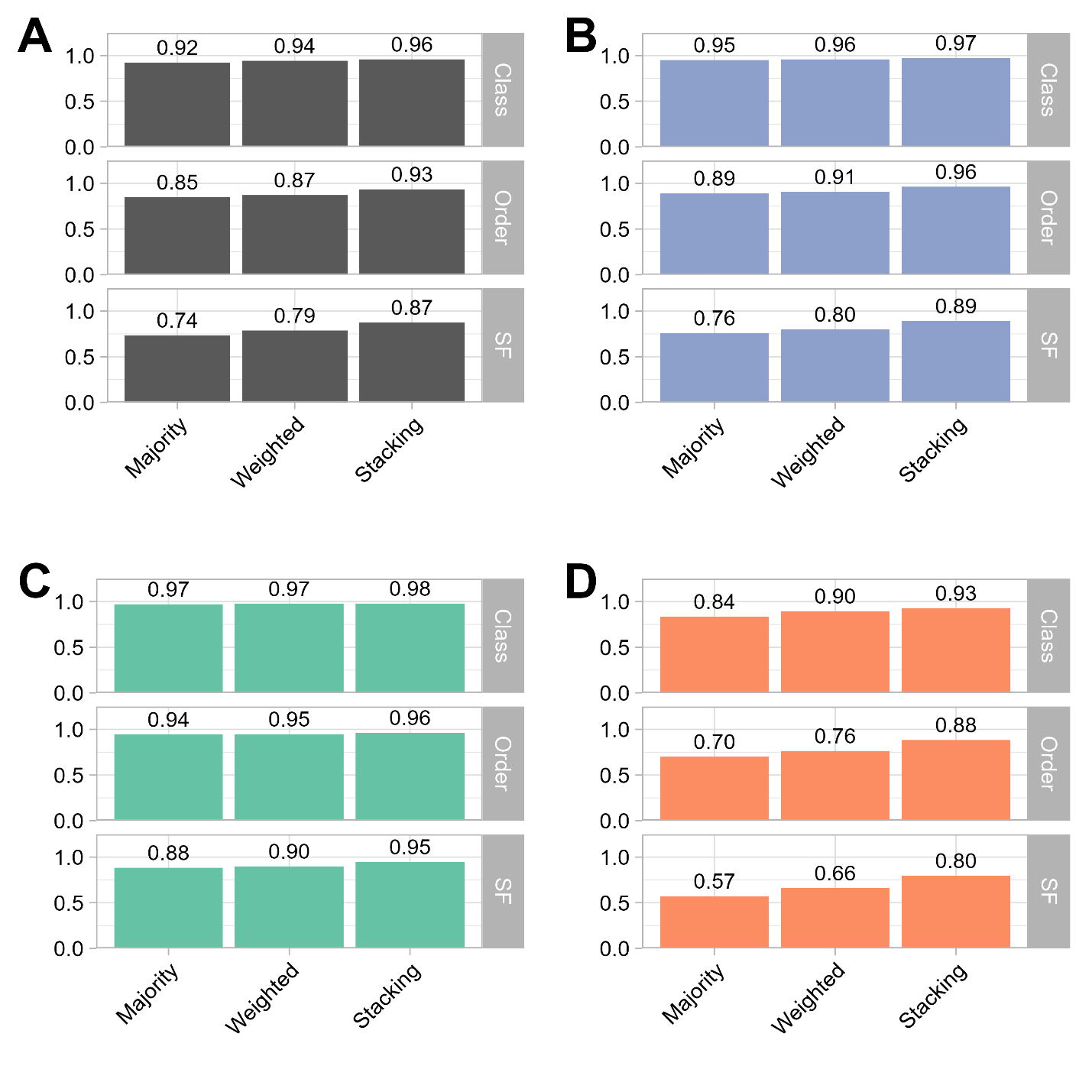

**Figure S6.** Voting systems performance using A) whole dataset, B) Animalia, C) Plantae and D) fungi.

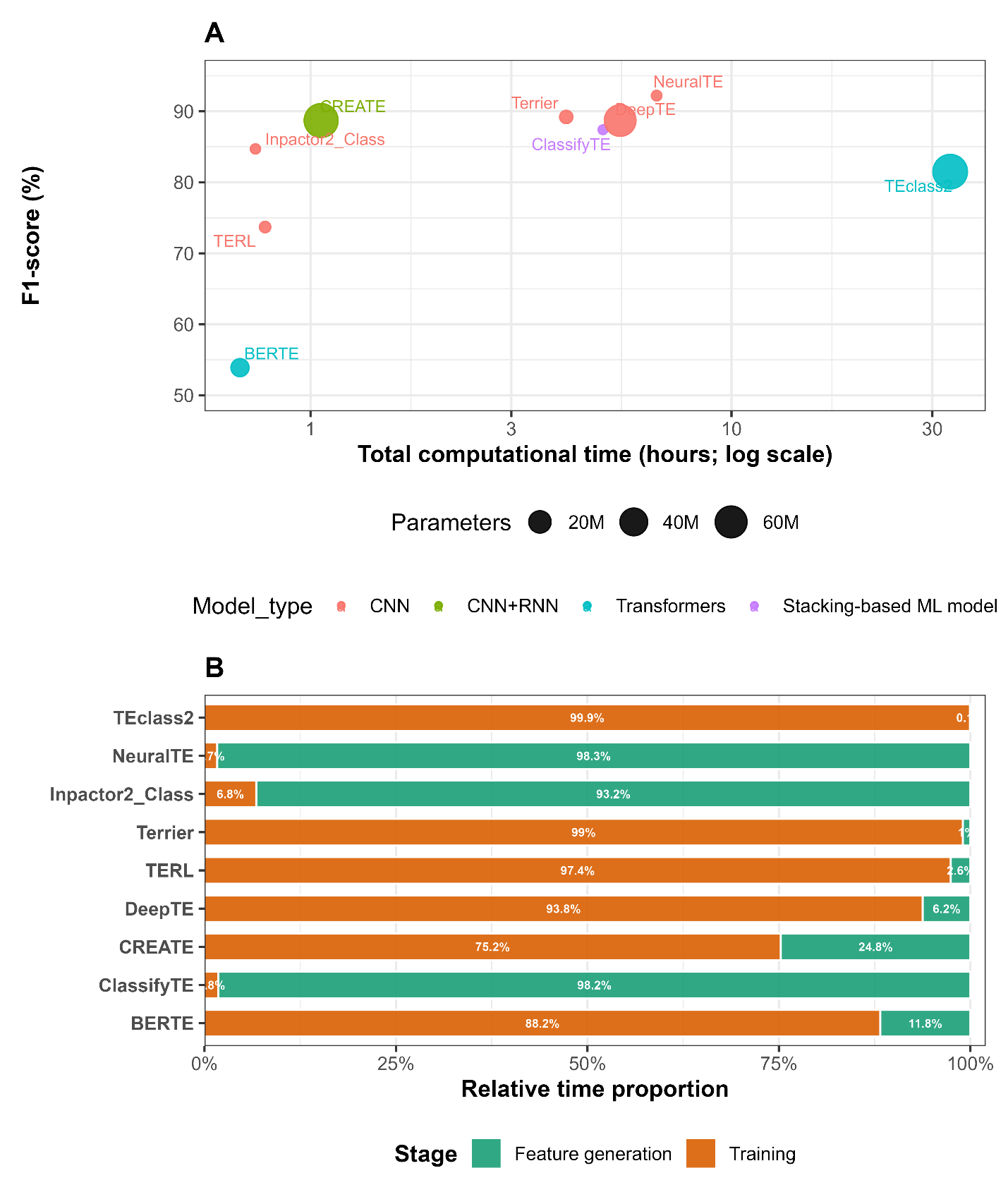

**Figure S7.** Computational performance trade-offs across TE classifiers. (A) Relationship between model performance (F1-score, y-axis), total runtime (x-axis, log scale), and model complexity (point size = number of trainable parameters; color = model type). Models located toward the upper-left region achieve higher accuracy with lower computational cost. (B) Relative contribution of feature generation (green) and model training (orange) to total runtime, expressed as percentages of the total execution time. The execution times can be consulted at Table S6.

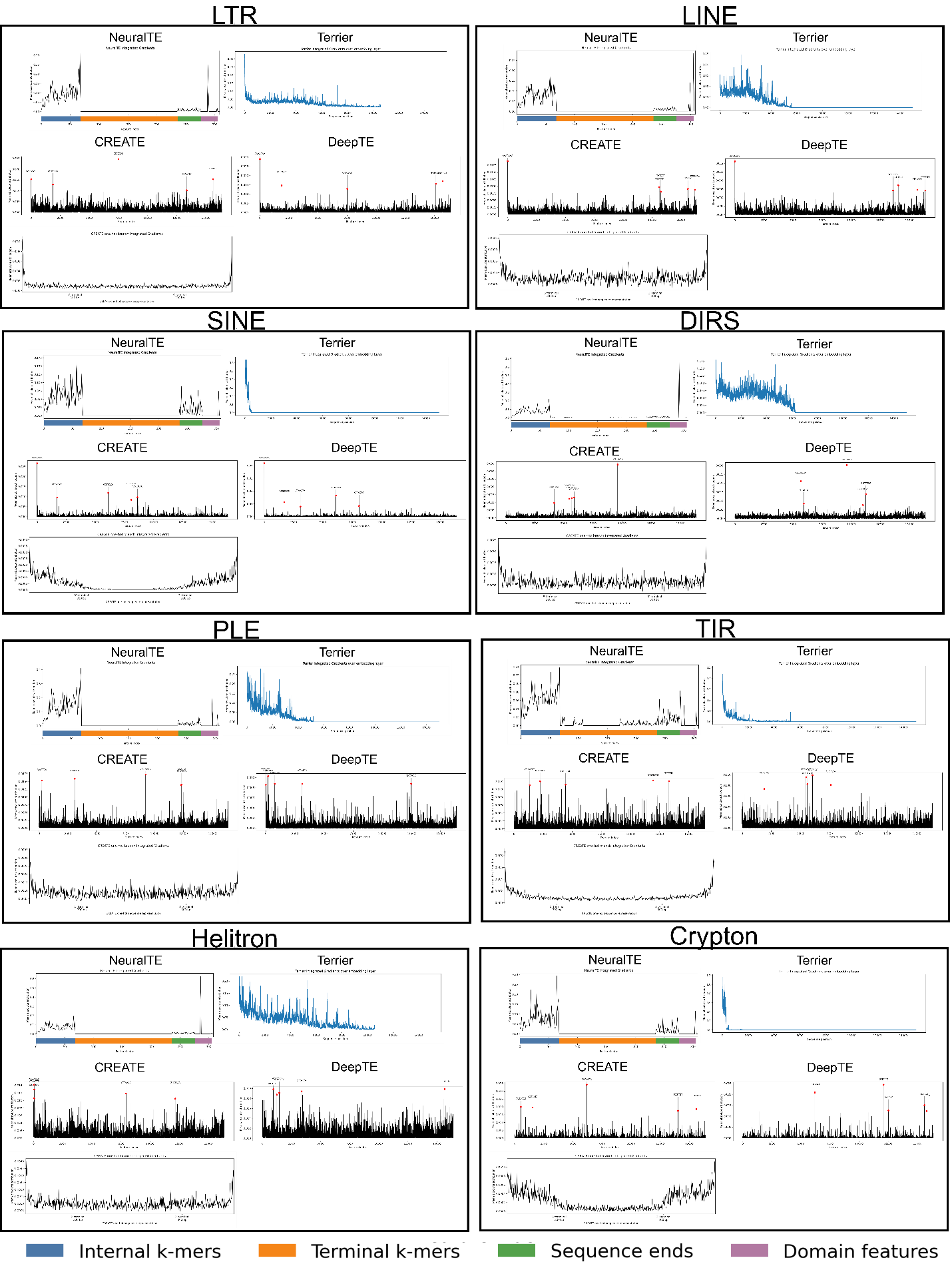

**Figure S8.** Explainability analyses of the four best-performing models: NeuralTE, Terrier, CREATE, and DeepTE. Each panel shows the features that contributed most to the classification of 20 randomly selected transposable elements representing different TE orders. Red points in CREATE and DeepTE *k*-mer panels highlight the five more contributing features.

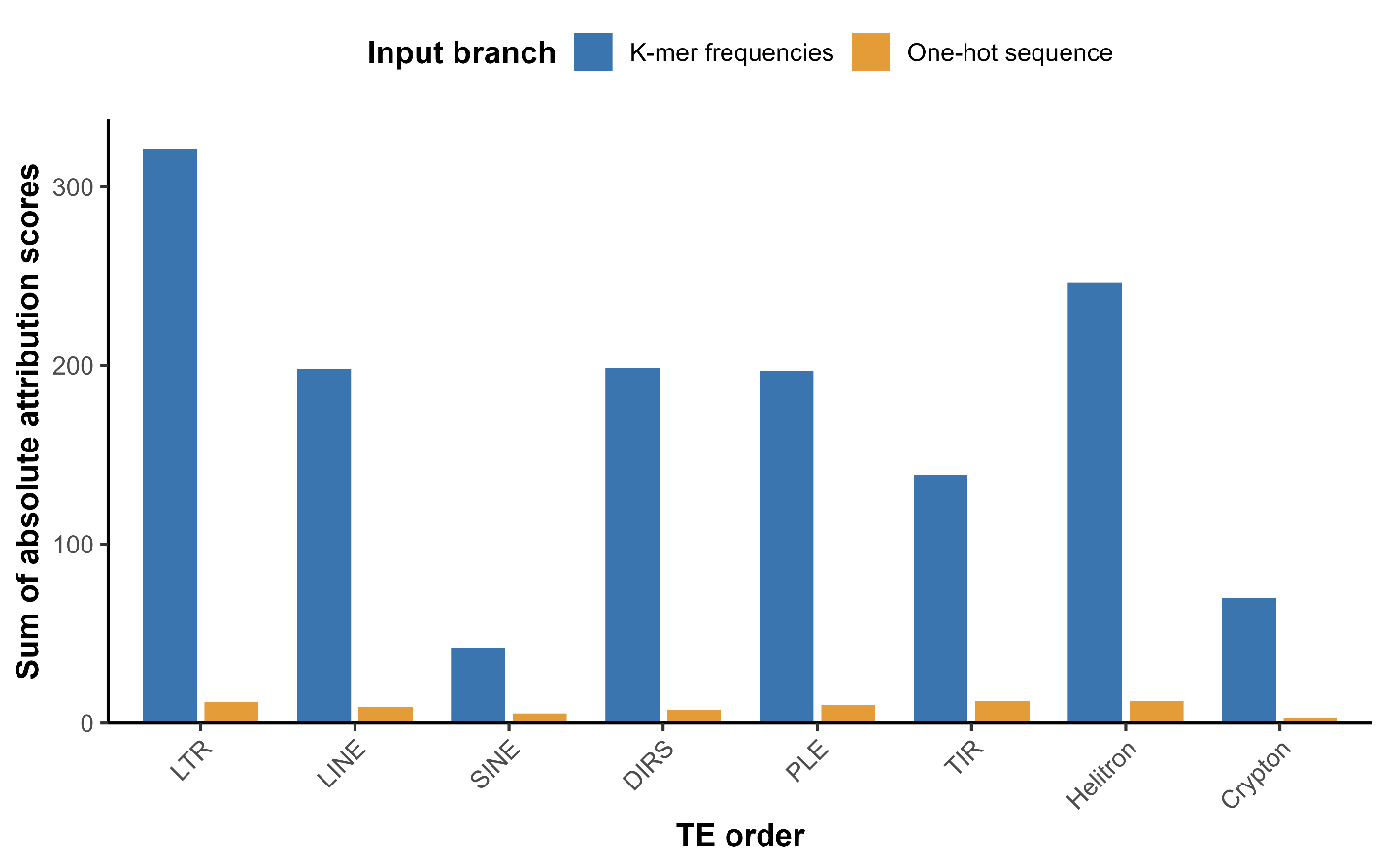

**Figure S9.** Relative contribution of the two input branches of CREATE to TE classification across TE orders. Bars represent the sum of the absolute attribution scores obtained by Integrated Gradients for the *k*-mer frequencies and one-hot sequence branches across the eight TE orders presented in the training dataset. Attribution values were summed across all features within each branch for each TE order.

**Supplementary Information**

**TE annotation of *C. glabrata* isolates using the PanTEon Database**

To demonstrate a practical application of the PanTEon Database to genome annotation, we annotated TEs in 12 *C. glabrata* isolates sequenced by Marcet-Houben and co-workers (Marcet-Houben et al., 2022). Because the database did not contain a species-specific TE library for *C. glabrata*, we extracted two reference libraries representing different taxonomic levels: the phylum Ascomycota and the family Debaryomycetaceae. We generated these libraries using “python3 PanTEon.py library --taxon LINEAGE”, replacing LINEAGE with Ascomycota or debaryomycetaceae in separate runs. Both libraries were exported in fasta format compatible with RepeatMasker (Smit et al., 2015). Genome assemblies were downloaded from NCBI BioProject PRJNA717653. RepeatMasker v4.2.3 was then run separately with each library, using default parameters except for -lib, which specified the corresponding PanTEon-derived library.

Using the Ascomycota-derived PanTEon library increased RepeatMasker annotation coverage in all 12 *C. glabrata* strains compared with the Debaryomycetaceae-derived library (Figure S1). Mean genome coverage increased from 0.122% to 0.383%, corresponding to a gain of 0.261 percentage points and a relative increase of 214.8%. Across the 12 assemblies, inferred TE copy counts increased from 563 to 4,060, representing a net gain of 3,497 annotations and 7.21 times as many inferred copies. Individual strains gained 278–319 copy annotations. The largest contributions came from LTR retrotransposons, whose counts increased from 110 to 1,586 (+1,476), and SINEs, which increased from 452 to 1,446 (+994). The Ascomycota library also yielded 606 LINE and 146 Helitron annotations, neither of which was detected using the Debaryomycetaceae library, while TIR annotations increased from one to 276. LTR retrotransposons accounted for 55.4% of the net increase in annotated base pairs, indicating that the broader library expanded both the coverage and diversity of putative TE annotations.

Comparisons among the eight sampled clades revealed differences primarily associated with LTR coverage. With the Ascomycota library, mean genome coverage was lowest in clade I (0.342%), intermediate in clade IIa (0.376%), and higher in clade VII (0.400%). Mean LTR coverage increased from 0.203% in clade I to 0.259% in clade VII, accounting for approximately 97.9% of their difference in total TE coverage. SINE coverage was comparatively uniform across all strains, ranging from 0.080% to 0.082%. M12, the sampled representative of clade III, exhibited the highest total coverage: 0.453% with the Ascomycota library versus 0.130% with the Debaryomycetaceae library, accompanied by an increase from 51 to 355 inferred copies. CBS 138, representing clade V, had the highest copy count and largest numerical gain, increasing from 47 to 366 copies (+319), although its coverage (0.412%) remained below that of M12. Thus, the strain with the greatest annotated coverage did not have the most inferred copies, reflecting differences in the amount of sequence assigned per copy. These clade comparisons remain descriptive because five of the eight clades were represented by a single strain.

**Architectonic benchmarking analysis**

Comparing the performance of different models within a common, standardized platform allowed us to investigate what differential factors may impact overall algorithm performance, thereby guiding future development efforts. Most tested algorithms relied on features generated directly from the DNA sequences, without any prior knowledge: one-hot and DAX (starting in 1 not in 0 as originally proposed (Yu et al., 2015) coding schemes, and *k*-mer frequency counts. These features are fast to retrieve and extract informative motifs that the model can learn to distinguish between classes (superfamilies in this study).

Nevertheless, in some cases the information contained in such features is insufficient for good performance due to the high heterogeneity at the nucleotide level characteristic to TEs. To overcome this, some researchers have applied more sophisticated ways to transform DNA sequences into numerical features through embeddings based on genomic language models. Even though this approach has obtained very promising results in other genomic tasks, we observed poor results in this study, probably because of the limited number of samples contained in the PanTEon Database. We hypothesize that once the current databases grow to have several hundreds of thousands of TE sequences, this approach will start to overcome other coding schemes.

One of the best performances obtained in our tests was obtained by calculating structural features in combination with *k-mer* counts. Characteristics like presence/absence of terminal repeats, and similarities with coding domains of known TEs are biologically meaningful information that the model could use to improve classification performance, albeit at the expense of increasing computational costs. Indeed, in NeuralTE, 98.3% of the runtime was used in generating features, and training was performed in the remaining (Figure S7B).

In terms of neural network architectures, CNN was the most popular option and generally resulted in high F1-scores. In contrast, the two Transformers-based architectures were the ones with the lowest scores, probably because of the reduced size of the training set relative to the complexity of DNA sequences. Transformer-based architectures generally achieved lower performance than several CNN- and feature-based approaches. While the relatively limited size of the training dataset compared with those typically used to train large language models likely contributed to this result, additional factors may also play an important role. First, TE classification is strongly influenced by local sequence motifs, structural signatures, and protein domains, features that can be effectively captured by convolutional architectures and handcrafted feature representations. Second, many transformer models operate on fixed-length token windows and may not fully exploit biologically relevant information distributed across long TE sequences. Finally, existing genomic foundation models are typically pretrained on general genomic sequences rather than curated TE collections, potentially limiting the transferability of their learned representations to TE classification tasks. Future work involving larger TE datasets, TE-specific pretraining strategies, and foundation models explicitly designed for repetitive DNA may help clarify the full potential of transformer architectures for this application. Interestingly, model size (measured as the number of training parameters) correlated with computing time but not with performance (Figure S7A). Terrier demonstrated a good balance between F1-score and computational time. Small computing times result from combining a simple feature extraction approach and a relatively small model size. The good performance was likely achieved thanks to the tool’s approach to compute training loss, which considers not only the predicted superfamily (as commonly done by other models), but also the hierarchical relation between orders and superfamilies. Although NeuralTE and Terrier obtained the best F1-Scores in our experiments, the way they create features or measure the training loss were designed to specifically classify TE superfamilies, so their usage may be limited for other classification tasks. In contrast, CREATE used only DNA sequence features, like one-hot coding and *k-mer* counts, which makes the algorithm more generalizable for other classification tasks.

The broader implications of PanTEon extend beyond classification accuracy. By revealing the taxonomic and structural signals learned by DL models, PanTEon exposes fundamental limitations in current TE ontologies and emphasizes the need for harmonized, machine-readable nomenclature. Its unified benchmarking environment provides a scalable and evolutionarily informed basis for TE annotation in large international genome consortia, where inconsistency in TE classification remains a major bottleneck for comparative genomics. At the same time, the biologically meaningful latent representations learned by PanTEon models open new avenues for evolutionary inference, including the discovery of novel TE families, reconstruction of deep homologies, and refined detection of horizontal transfer.

**Explainability analysis**

After finding the best performing AI architectures we were interested in going deeper and understanding how the models were learning the different used features, how they learned the TE biology and the differences between predicting sequences from different TE orders. So, we developed a model-specific interpretability pipeline based on Integrated Gradients (IG), a gradient-based attribution method widely used for explaining genomic sequence models (Majdandzic et al., 2023; Novakovsky et al., 2023; Sundararajan et al., 2017). We randomly selected 20 sequences from the eight TE orders presented in the training dataset (LTR, LINE, SINE, DIRS, PLE, TIR, Helitron, and Crypton) and used the four best performing architectures: NeuralTE, Terrier, CREATE and DeepTE (see Methodology for details).

NeuralTE calculates four different types of features: frequency of internal *k*-mers (*k*=1,2), frequency of terminal *k*-mers (*k*=1,2,3), sequence ends (coded through DAX scheme), and presence of coding domain. Interestingly, the IG analysis showed that internal *k*-mers contributed the most when classifying all the TE orders, while terminal *k-*mers were not adding any importance at all for classifying sequences form all the TE orders, except for TIR elements (Figure S8). This finding indicates that NeuralTE primarily relied on conserved sequence features located within internal regions rather than terminal structures (Majdandzic et al., 2023; Sundararajan et al., 2017). This observation is consistent with the fact that the internal regions of most autonomous TEs contain conserved protein-coding domains, such as reverse transcriptases, integrases, helicases, and transposases, which represent the primary evolutionary signatures used for TE classification (Orozco-Arias et al., 2019; Poulter & Butler, 2015; Thomas & Pritham, 2015; Wicker et al., 2007). In contrast, TIR elements constituted the only notable exception, as terminal *k*-mers received attribution scores, likely reflecting the presence of important motifs, which are defining structural features required for transposase recognition and transposition (Feschotte & Pritham, 2007; Wicker et al., 2007). Finally, the presence of TE-specific coding domains substantially contributed to accurate classification. For instance, in LTR retrotransposons, domains characteristic of the Copia and Gypsy superfamilies ranked among the 20 most important features, whereas domains associated with other TE orders contributed negligibly. A similar pattern was observed across the remaining TE orders, with attribution scores largely restricted to domains characteristic of the predicted order. Specifically, LINE elements were primarily supported by domains from the LINE, LINE/RTE, and LINE/L1 superfamilies; DIRS elements by DIRS-specific and unclassified domains; PLEs by PLE-specific and unclassified domains; TIR elements by Tc1-Mariner, hAT, PIF-Harbinger, and unclassified domains; Helitrons by Helitron, LINE, and unclassified domains; and Cryptons exclusively by unclassified domains. In contrast, the non-autonomous SINE elements received attribution almost exclusively from unclassified domains, consistent with their lack of protein-coding capacity.

Terrier followed a different strategy by representing sequences exclusively with DAX-encoded nucleotides, while reserving zero for padding and allowing input sequences of up to 15 kbp. Consequently, the attribution profiles varied substantially among TE orders. For LTR retrotransposons and TIR elements, the first 64 bp and 59 bp, respectively, exhibited markedly higher attribution scores than the remainder of the sequence (Figure S8). This pattern is consistent with the presence of characteristic terminal repeats and conserved sequence motifs at the 5′ ends of these elements. Surprisingly, the corresponding 3′ terminal repeats did not display comparable attribution scores. Instead, additional attribution peaks were observed around positions 3.8, 4.6, and 7.2 kbp in LTR retrotransposons and around 5.5 kbp in TIR elements. We hypothesize that these peaks correspond to the approximate positions of the opposite terminal boundaries in different superfamilies, although their reduced attribution scores are likely a consequence of sequence length heterogeneity and the extensive padding introduced to accommodate variable-length sequences. Terrier's encoding strategy also provides an additional advantage for TE classification by implicitly incorporating sequence length information. Because sequences are padded with zeros at the 3′ end to a fixed length of 15 kbp, the model can infer the approximate length of each element as an informative feature. This effect is particularly evident for LINEs, where nucleotides beyond approximately 6 kbp showed no contribution to the predictions, and for SINEs, where attribution scores dropped to zero beyond approximately 500 bp.

CREATE employed a dual-branch architecture that processed two complementary feature representations: 7-mer frequency vectors and one-hot encoded representations of the 5′ and 3′ terminal 300 bp of each sequence. Despite this complementary design, the *k*-mer frequency branch consistently contributed more to the final predictions than the one-hot sequence branch across all TE orders, with summed attribution scores ranging from 7.6-fold higher for SINE elements to 26-fold higher for LTR retrotransposons (Figure S9 and Table S20). These results indicate that global sequence composition captured by *k*-mer frequencies provides substantially greater discriminatory power than terminal nucleotide patterns alone. This observation is consistent with previous studies showing that *k*-mer frequency features outperform alternative sequence representations for the classification of LTR retrotransposons (Orozco-Arias et al., 2021). Interestingly, the attribution pattern observed for CREATE's one-hot encoded branch closely resembled those obtained with NeuralTE and Terrier, despite these models relying on different sequence representations. In all three models, the highest attribution scores were concentrated at the 5′ and 3′ termini of LTR retrotransposons and TIR elements. Elevated attribution was also observed for DIRS and Helitrons, which possess characteristic terminal repeat structures, although the signal was substantially weaker than that of LTR retrotransposons and TIR elements.

When analyzing the CREATE’s *k-*mer branch, the most influential 7-mers were highly order-specific, with only the homopolymeric motif AAAAAAA appearing among the top contributors for multiple retrotransposon orders (LTR, LINE, and SINE). In contrast, the remaining highly attributed *k*-mers differed markedly among TE orders, suggesting that the classifier learned characteristic sequence signatures rather than relying on universal motifs (Figure S8). This behavior was particularly evident for TIR elements and Cryptons, whose 5 top-ranked *k*-mers showed little overlap with those of other orders, supporting the notion that the *k*-mer representation captures lineage-specific sequence composition beyond conserved terminal structures.

DeepTE exhibited a similar attribution pattern (Figure S8), with the highest contributions concentrated in internal *k*-mers and only minor contributions from terminal regions. As observed for CREATE, the most informative 7-mers were largely specific to individual TE orders, although a greater overlap between related retrotransposon orders was apparent, with motifs such as AAAAAAA and ACGGTCC contributing to multiple classes. This suggests that DeepTE exploits both order-specific sequence signatures and compositional similarities among evolutionarily related TE groups. Notably, the 5 top-ranked *k*-mers identified by both models rarely corresponded to known terminal repeat motifs, reinforcing the conclusion that internal sequence composition provides substantially greater discriminatory power than terminal sequence patterns for TE order classification.

Although these analyses are not exhaustive, they demonstrate the potential of explainable AI to uncover the biological principles learned by ML/DL models for TE classification. Future studies combining multiple attribution methods, larger and taxonomically broader TE datasets, and foundation models specifically trained on repetitive DNA will enable a more comprehensive characterization of the sequence features underlying TE evolution and classification, ultimately facilitating the development of more accurate and biologically interpretable AI models.
